# Spatial multi-omic profiling of non-autonomous transcriptomic and epigenomic responses to genetic perturbation

**DOI:** 10.64898/2026.11.12.766149

**Authors:** Javid Ghaemmaghami, Kun H. Lee, Shean Jiang, Yalei Zhao, Gaurav Gadhvi, Lu Lu, Ming-Ching Wen, Yixin Wang, Liu Liu, Dylan Cable, Joshua D. Welch

## Abstract

Multicellular organisms develop and maintain differentiated cell states through a combination of cell-autonomous and non-autonomous regulatory mechanisms, including signal transduction pathways mediated by ligands, receptors, and kinases. Genetic perturbation through CRISPR pooled screening is widely used to study cell-autonomous regulatory mechanisms, but the challenges of cellular-resolution *in situ* measurement have limited the study of non-autonomous regulatory mechanisms. We developed Perturb-Tags, a spatial multi-omic technique for pooled CRISPR screening that tags nuclei with spatial barcodes, allowing joint measurement of single-nucleus gene expression, CRISPR guide RNAs, chromatin accessibility, and spatial positions within intact tissue. Perturb-Tags achieves cellular-resolution, unbiased gene expression and multi-omic profiling, and higher sensitivity than previous spatial perturbation approaches. We used Perturb-Tags to systematically perturb ligands, receptors, and all expressed kinases in human embryoid bodies. These experiments revealed extensive non-autonomous gene expression changes after perturbation, including distance-dependent loss of signaling pathway activity upon ligand knockdown, compensatory upregulation of alternative signaling pathways, and responses to receptor and kinase knockdown that vary with cell microenvironment. We also identified changes in the spatial organization of cells with ligand and kinase knockdowns. Finally, we trained a graph neural network to model the combined effects of genetic perturbations on gene expression and chromatin accessibility across an entire tissue. Our work provides a powerful experimental and computational platform for mechanistic studies of non-autonomous regulatory mechanisms across healthy and diseased tissues.

## Main

The development and function of multicellular tissues depend on both cell-autonomous regulatory programs in individual cells and non-autonomous signals exchanged among neighboring cells^1,2^. Non-autonomous interactions are particularly important because perturbations in one cell can propagate through signaling networks to alter the state, fate and function of surrounding cells, thereby shaping collective tissue behavior^1,3^. Such effects are central to developmental patterning and tissue homeostasis and can amplify or redirect pathological processes during disease progression, making it essential to measure not only the response of the perturbed cell but also its consequences across the surrounding cellular neighborhood^2,3^. Ligands, receptors, intracellular kinases, and transcription factors form signal transduction cascades that coordinate processes including embryonic patterning, lineage specification, immune responses, and tissue homeostasis^1,2,4^. Dysregulation of signal transduction pathways contributes to diverse diseases^2,4–6^. Pooled CRISPR screening approaches such as Perturb-seq and CROP-seq have enabled systematic measurement of cell-autonomous perturbation responses by linking guide identity to transcriptome-wide expression in individual cells^7–9^. However, because these methods require tissue dissociation, they lose the spatial relationships needed to determine how perturbations influence neighboring cells and cell non-autonomous interactions^10–12^. This limitation is particularly important for cell–cell communication pathways, in which perturbing a ligand, receptor, or intracellular signaling component can alter molecular states beyond the cell carrying the perturbation itself, with effects that depend on distance, cell identity, and local signaling context^1,2,11^. Recent methods combining pooled perturbation with spatial profiling have begun to address this problem^10–13^, but targeted imaging approaches are limited to preselected transcripts, whereas spot-based sequencing methods often have lower sensitivity and can mix material from multiple neighboring cells, complicating assignment of perturbations and responses to individual cells^11–13^. A method for unbiased joint measurement of perturbation identity, transcriptome-wide expression, chromatin accessibility, and spatial position in the same cell would therefore enable direct analysis of both cell-autonomous and non-autonomous perturbation effects.

Here we developed Perturb-Tags, a spatial multi-omic platform for pooled CRISPR screening that spatially barcodes nuclei before performing ex situ sequencing on a standard single-cell platform. Perturb-Tags jointly measures spatial coordinates, CRISPR guide RNAs (gRNAs), and transcriptome-wide gene expression from the same nucleus, with optional simultaneous measurement of chromatin accessibility. Thus, Perturb-Tags combines cellular-resolution spatial information with the sensitivity and breadth of sequencing-based single-nucleus multi-omic profiling. We applied Perturb-Tags to human embryoid bodies carrying CRISPR interference libraries targeting ligands and receptors involved in developmental signaling and, separately, a large-scale library targeting expressed kinases. These screens interrogate complementary levels of signaling, from extracellular signal production and reception to intracellular pathways that interpret local microenvironmental inputs. We further developed a graph neural network model that integrates perturbation identity, molecular state, spatial position, and neighboring cells to model how genetic perturbations propagate through tissue and alter gene expression and chromatin accessibility. Together, Perturb-Tags and this computational framework reveal how perturbations across multiple levels of signaling propagate through spatial cellular neighborhoods, while establishing a broadly applicable platform for dissecting multicellular regulation in development, homeostasis, and disease.

### Perturb-Tags measures gene expression, chromatin accessibility, and CRISPR guide RNAs in spatially barcoded nuclei

Spatial barcoding methods such as Slide-tags combine the molecular sensitivity of sequencing-based single-cell assays with cellular-resolution spatial information^14^. We reasoned that this strategy could be extended to capture CRISPR gRNAs, thereby linking perturbation identity to transcriptomic and epigenomic responses while preserving the position of each nucleus within the original tissue. We therefore developed Perturb-Tags, a spatial multi-omic pooled-screening method that jointly measures gene expression, CRISPR single gRNAs (sgRNAs), spatial coordinates, and, optionally, chromatin accessibility in the same nucleus (**Fig. 1a**). Perturb-Tags uses the commercially available Trekker spatial-barcoding assay and 10x Genomics Chromium platform, facilitating implementation using broadly accessible reagents and instrumentation^14,15^.

**Figure 1:**
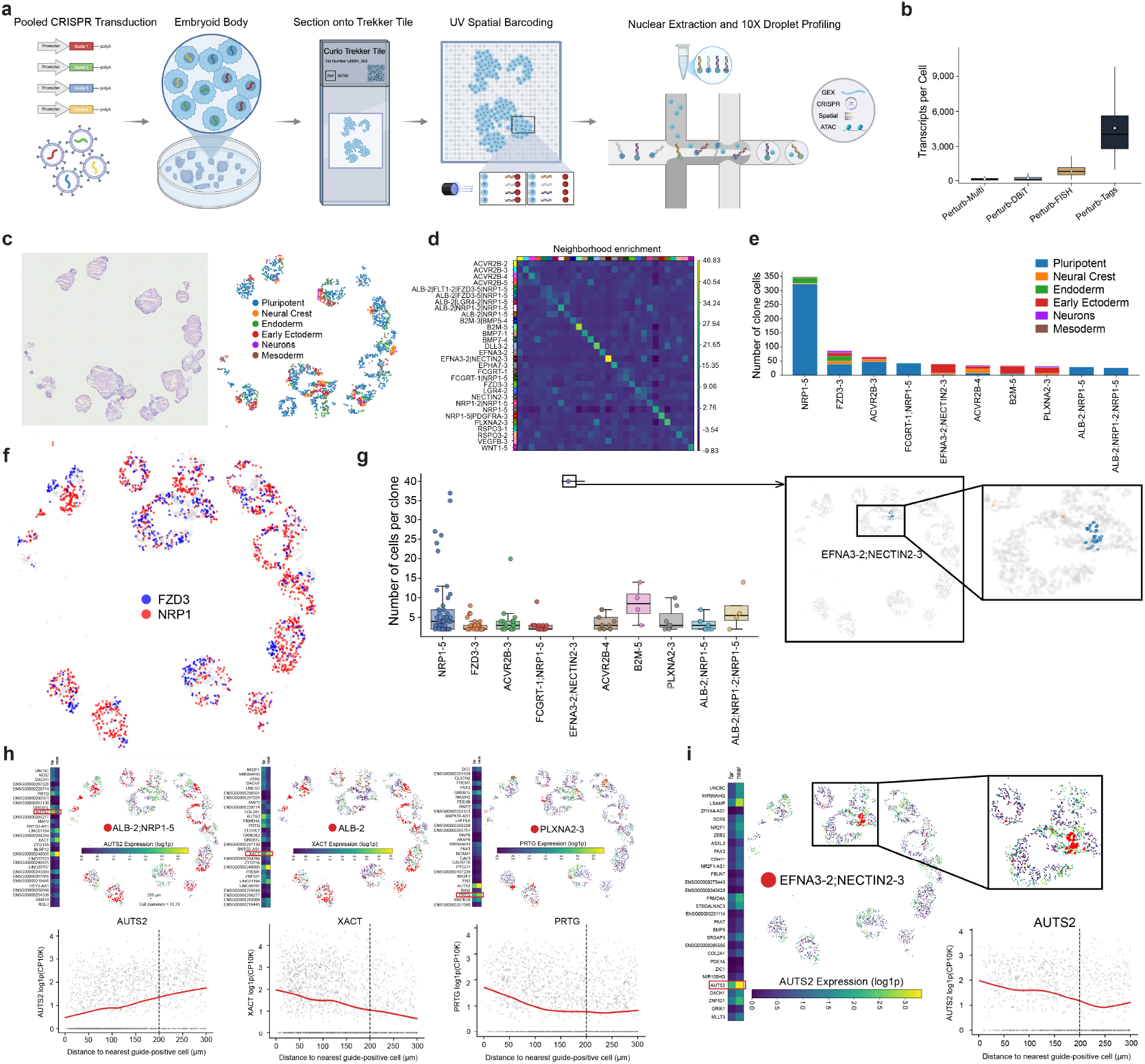
Perturb-Tags enables spatial multi-omic profiling of non-autonomous response to perturbation. **a**, Schematic of the Perturb-Tags workflow. Cells are transduced with a pooled CRISPR sgRNA library and differentiated as embryoid bodies (EBs). Multiple EBs are cryosectioned onto a Trekker tile and subjected to UV illumination to release photocleavable oligonucleotides carrying spatial barcodes, which are transferred to nuclei within the tissue section. Following nuclear extraction, nuclei are profiled using the 10x Genomics Chromium platform. Spatial barcodes, transcripts and sgRNAs are captured from the same nucleus, with optional simultaneous measurement of chromatin accessibility. **b**, Comparison of number of transcripts detected per cell or nucleus using Perturb-Tags and previously reported spatial perturbation-profiling approaches. **c**, H&E stain for adjacent tissue section (left) and spatial coordinates of Perturb-Tags nucleus locations colored by annotated cell type (right). We collected and analyzed Perturb-Tags data on EBs that had 24 expressed ligands and receptors knocked down. **d**, Neighborhood enrichment analysis of cells grouped by detected sgRNA identity. Rows and columns denote sgRNAs or sgRNA combinations, and color indicates the enrichment of each pair within local spatial neighborhoods. Enrichment along the diagonal indicates preferential spatial proximity between cells carrying the same sgRNA or sgRNA combination. **e**, Number and cell-type composition of cells assigned to spatial clones for selected sgRNAs and sgRNA combinations. Colors denote annotated cell types. **f**, Representative spatial distribution of cells carrying FZD3 or NRP1 perturbations across EBs, illustrating spatially restricted patches of cells sharing perturbation identities. **g**, Distribution of clone sizes for selected sgRNAs and sgRNA combinations, defined by spatially connected cells carrying the same complement of detected sgRNAs. Right, representative EFNA3-2;NECTIN2-3 clone and magnified view showing a spatially localized group of cells sharing the perturbation. **h**, Representative examples of distance-dependent receiver-cell gene expression surrounding perturbation neighborhoods. Bottom, scatter plots of receiver-cell expression as a function of distance to the nearest cell carrying the indicated perturbation; red lines are LOESS smooths indicating the distance-dependent expression trend. Scale bar, 200 µm. **i**, Spatial distribution of an EFNA3-2;NECTIN2-3 perturbation neighborhood and AUTS2 expression in surrounding receiver cells. Bottom right, receiver-cell AUTS2 expression as a function of distance to the nearest EFNA3-2;NECTIN2-3 cell.

We applied Perturb-Tags to human embryoid bodies (EBs), a tractable model of early development that is widely used for genetic studies and as a starting point for organoid differentiation protocols^16^. We transduced human induced pluripotent stem cells stably expressing dCas9-KRAB with a pooled lentiviral library in which each vector encoded a distinct CRISPR sgRNA. Following puromycin selection, we made EBs and grew them for two weeks, allowing perturbed cells to differentiate and proliferate within a spatially organized multicellular environment. To increase scalability and control technical variation, we performed Perturb-Tags on many EBs within a single experiment: we embedded and flash-froze multiple EBs within a single block and cryosectioned the block onto a bead-coated Trekker slide, so that each cryosection contained multiple EBs (**Fig. 1a**). Each bead on the slide carries oligonucleotides containing a spatial barcode connected through a photocleavable linker. Ultraviolet illumination releases these oligonucleotides from the beads and transfers the spatial barcodes to nuclei within the overlying tissue section. Because the position of each spatial barcode is known, the barcode sequence associated with a particular nucleus indicates its original coordinates within the tissue. Following spatial barcoding, we extracted the nuclei and loaded them onto the 10x Genomics Chromium system for droplet-based single-nucleus profiling. Transcripts, CRISPR sgRNAs, and nucleus-associated spatial barcodes are then captured within a single droplet, linking gene expression and perturbation identity to the position of the same nucleus in the tissue. To extend Perturb-Tags to joint transcriptomic and epigenomic profiling, we used a modified 10x Multiome workflow by adding a custom reverse-transcription primer and splint oligonucleotide that enabled sgRNA capture alongside RNA and transposase-accessible chromatin. Perturb-Tags can therefore generate a four-component readout comprising spatial position, gene expression, sgRNA identity, and chromatin accessibility from individual nuclei.

Perturb-Tags recovered transcripts and sgRNAs with high sensitivity. Transcript detection was comparable to that reported for conventional single-nucleus RNA sequencing of human EBs, indicating that spatial barcoding did not compromise transcriptome complexity. Perturb-Tags also detected substantially more transcripts per nucleus than previous spatial perturbation approaches, while retaining cellular resolution and unbiased transcriptome-wide coverage (**Fig. 1b**).

### Clonal expansion of perturbed cells reveals distance-dependent changes in ligand–receptor signaling

Perturb-Tags recovered transcripts and sgRNAs with high sensitivity. Transcript detection was comparable to that reported for non-spatial single-nucleus RNA sequencing of human EBs^17^, indicating that spatial barcoding did not compromise transcriptome complexity. Perturb-Tags also detected substantially more transcripts per nucleus than previously reported spatial perturbation approaches, while retaining cellular resolution and unbiased transcriptome-wide coverage (**Fig. 1b**).

We first used Perturb-Tags to screen the effects of knocking down 26 ligands and receptors involved in important developmental signaling interactions, including BMP, WNT, Notch, ephrin, and semaphorin signaling pathways (**Fig. 1c**). In our first experiment, we recovered 4,115 cells with a median of 4,084 transcripts per cell, median of 2,600 genes per cell, and 65.7% of cells with detectable guide RNAs. Because we grew the embryoid bodies for two weeks after transduction, we expected that dividing cells may have propagated the integrated sgRNA vector to daughter cells, creating spatially adjacent patches of clonal cells. Consistent with the expectation, we found that specific guides and guide combinations are enriched in the spatial neighborhood of cells containing the same guide (**Fig. 1d**). To further investigate this phenomenon, we built a spatial graph connecting nearby cells with the exact same complement of detected sgRNAs and identified connected components containing spatial clusters indicating potential clonal expansion (**Methods**). We observed that detected sgRNAs were frequently organized into spatially restricted patches rather than being randomly distributed throughout each embryoid body (**Fig. 1e-g**). The sizes of the spatial patches varied across guide combinations (**Fig. 1 e,g**). This heterogeneity may reflect differences in developmental timing, stochastic differences in clonal expansion, or perturbation-specific effects on cell proliferation or survival. Clone composition was also strongly associated with cell identity. For example, cells with NRP1 knockdown (KD) were predominantly pluripotent, whereas cells with joint EFNA3 and NECTIN2 KDs or B2M KDs were largely composed of Early Ectoderm cells (**Fig. 1e**). The spatial adjacency of cells sharing the same sgRNAs conveniently enables detecting coordinated non-autonomous perturbation effects in the local microenvironment.

We next performed a simple test to identify gene expression differences between cells near and far from cells with a particular perturbation. For each gene targeted in the screen, we excluded cells carrying that specific perturbation and defined the remaining cells as receiver cells. We then classified receiver cells as near or far from cells carrying the perturbation of interest and tested for differentially expressed genes between the two groups. Single and multiple-perturbation gene combinations exhibited distance-dependent expression patterns, demonstrating that local perturbation density was associated with transcriptional changes extending beyond the directly perturbed cells (**Fig. 1h,i**). For example, AUTS2 expression increased with distance from cells carrying ALB-2;NRP1-5 (**Fig. 1h**), but decreased with distance from cells carrying EFNA3-2;NECTIN2-3 (**Fig. 1i**). Similarly, XACT and PRTG expression were highest near ALB-2 and PLXNA2-3 cells, respectively, and decreased with distance. Together, these results indicate detectable distance-dependent expression differences near spatial clusters of cells sharing a common genetic perturbation.

Given the distance-dependent transcriptional changes around perturbation clusters, we next investigated whether these perturbations also altered local cell–cell signaling. Using CytoSignal^18^, we quantified ligand–receptor (LR) scores in receiver cells as a function of distance from the corresponding knockdown clone (again removing cells carrying the specific guide being analyzed). We examined both direct interactions involving the targeted ligand and compensatory interactions involving related or alternative signaling components (**Fig. 2a**,**b**). Spatial maps showed localized direct LR changes following BMP7 and ALB KD (**Fig. 2c**) and compensatory changes following JAG1 and NECTIN3 KD (**Fig. 2d**). Across the screen (**Extended Data Figs.1-3**), 34 direct LR interactions showed significant distance-dependent responses (**Fig. 2e**), including reduced EFNA3–EPHA1 signaling near EFNA3 KD cells (**Fig. 2f**). We also identified 25 compensatory LR interactions (**Fig. 2g**), including elevated DLK1–NOTCH1 signaling near DLL3 KD cells that decreased with distance (**Fig. 2h**). Among the direct LR interactions showing significant distance-dependent effects, the majority showed decreased local signaling near knockdown cells; the majority of significant compensatory interactions showed increased local signaling near knockdown cells. Together, these results support both direct and compensatory cell non-autonomous changes in signaling around spatially clustered perturbations.

**Figure 2:**
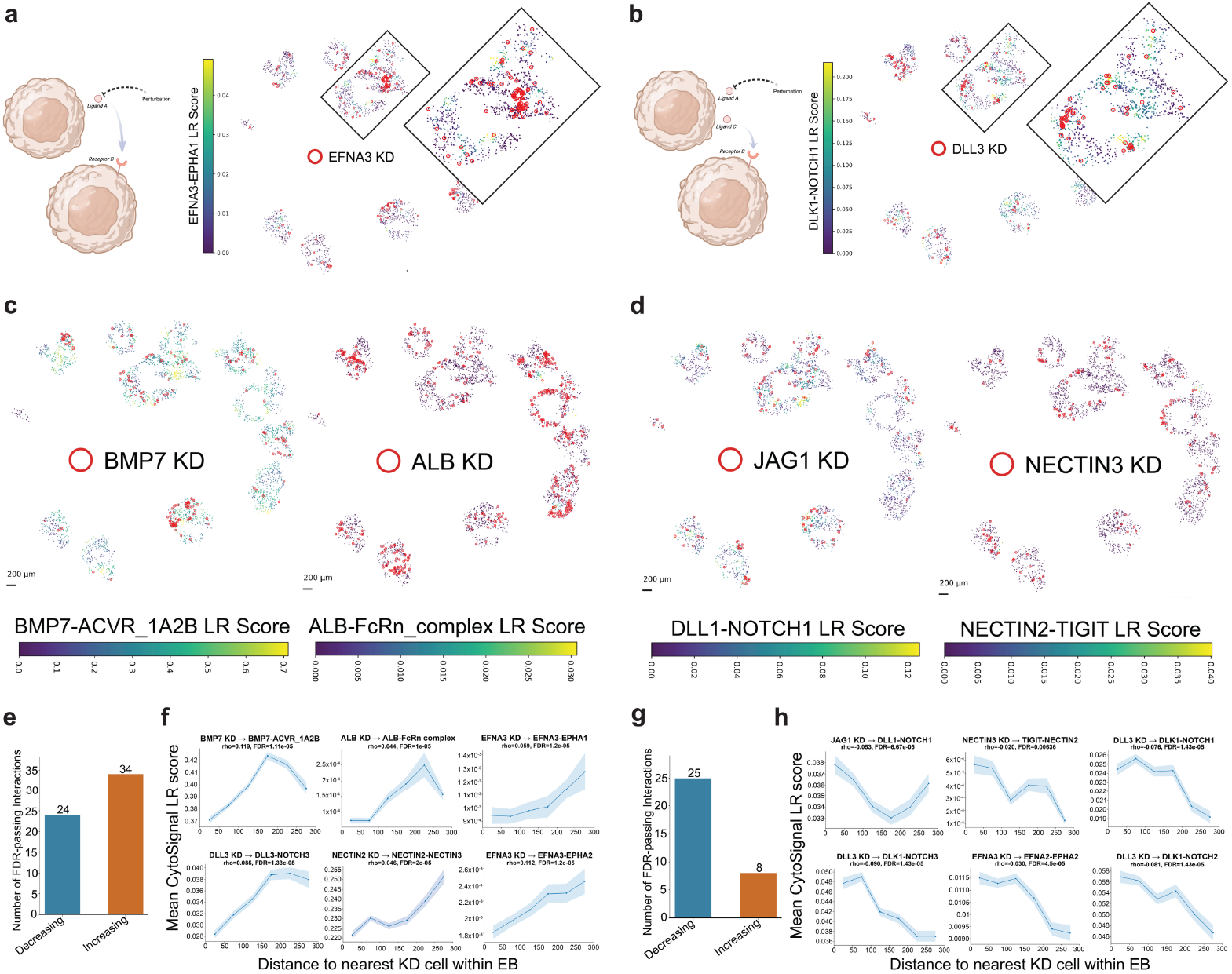
Perturb-Tags reveals direct and compensatory changes in ligand–receptor signaling near ligand knockdown cells. **a**, Schematic of a direct signaling response and representative spatial map of the EFNA3–EPHA1 ligand–receptor (LR) score following EFNA3 knockdown (KD). Receiver cells are colored by LR score; red open circles indicate cells carrying the corresponding perturbation. **b**, Schematic of a compensatory signaling response and representative DLK1–NOTCH1 LR scores surrounding DLL3 KD cells. **c**, Representative direct LR responses following BMP7 and ALB KD, showing spatial variation in BMP7–ACVR1A2B and ALB–FcRn complex scores, respectively. **d**, Representative compensatory LR responses following JAG1 and NECTIN3 KD, showing spatial variation in DLL1–NOTCH1 and NECTIN2–TIGIT signaling, respectively. **e**, Number of direct LR interactions passing FDR correction that decreased or increased with distance from the corresponding KD cells. **f**, Representative direct LR interactions showing mean CytoSignal LR score as a function of distance to the nearest corresponding KD cell within each embryoid body (EB). **g**, Number of compensatory LR interactions passing FDR correction that decreased or increased with distance from KD cells. **h**, Representative compensatory LR interactions showing mean CytoSignal LR score as a function of distance from the nearest KD cell within each EB. Direct interactions were defined as LR pairs containing the ligand targeted by the perturbation, whereas compensatory interactions involved related or alternative ligand–receptor pairs whose signaling changed following perturbation of another pathway component.

### Distance-dependent transcriptomic and epigenomic changes near cells with ligand perturbations

To determine whether the non-autonomous transcriptional responses observed in the initial Perturb-Tags experiment were accompanied by changes in epigenomic regulatory state, we generated a second dataset using the 10X Multiome assay, enabling joint measurement of gene expression and chromatin accessibility while preserving spatial organization (**Fig. 3a,b**). A single Trekker slide was distributed across three 10x Chromium lanes, yielding approximately 26,000 nuclei, of which 46% could be assigned one or more CRISPR sgRNAs. The assay recovered high-complexity RNA profiles and a median of approximately 15,000 ATAC fragments per nucleus, demonstrating that spatial barcoding and sgRNA capture were compatible with joint transcriptomic and chromatin-accessibility profiling. Importantly, this approach maintained robust transcript UMI capture without compromising RNA sensitivity (**Extended Data Fig. 4a**). We performed two different kinds of statistical tests to identify chromatin accessibility features that showed a significant change as a function of distance from cells with ligand knockdowns. We employed a spatial permutation test and Gaussian process regression to detect ATAC peaks, fixed-size genomic bins, ATAC gene activity scores, and chromVAR motif scores that increased or decreased with distance from ligand knockdown cells (**Fig. 3c, Extended Data Figs. 5 and 6**). We similarly tested how transcriptional effects depended on the local ligand microenvironment across different spatial radii, revealing pathway-specific responses that peaked at distinct spatial scales (**Fig. 3d**). We similarly detected genes with significant distance-dependent expression trends. Using a hierarchical Gaussian process model^19^, we clustered these significant genes by their spatial trends to identify coordinated gene-expression programs with distinct distance-dependent trajectories (**Fig. 3e**).

**Figure 3:**
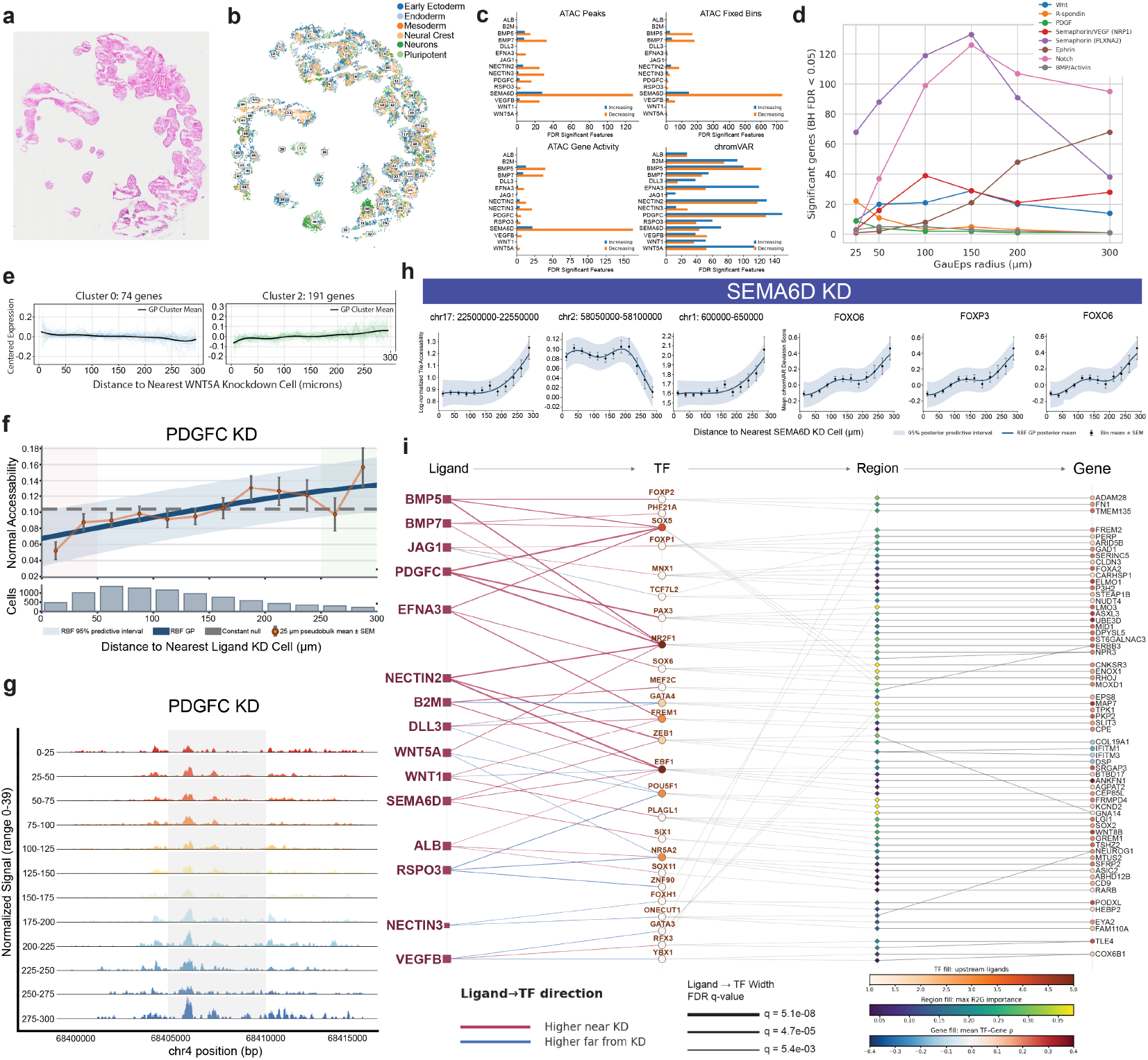
Distance-dependent gene expression and chromatin accessibility changes near cells with ligand and receptor perturbations. **a**, H&E staining of the tissue profiled by Perturb-Tags multiome. **b**, Spatial distribution of recovered nuclei colored by annotated cell type. **c**, Number of significantly distance-dependent features following each perturbation across ATAC peaks, fixed ATAC bins, ATAC gene activity and chromVAR motif activity, separated by increasing and decreasing responses. **d**, Number of significantly distance-dependent genes (BH FDR < 0.05) identified across signaling pathways as a function of the spatial scale used for Gaussian-process modeling. **e**, Representative clusters of genes exhibiting decreasing or increasing expression with distance from WNT5A KD cells. **f**, Representative Gaussian-process fit showing distance-dependent chromatin accessibility following PDGFC KD. Points indicate 25-μm pseudobulk means ± s.e.m.; the fitted spatial model and 95% posterior predictive interval are shown. **g**, ATAC fragment coverage across successive 25-μm distance bins for the representative PDGFC KD shown in the upper panel. **h**, Representative distance-dependent changes in regional chromatin accessibility and chromVAR transcription-factor motif activity following SEMA6D KD. **i**, Integrated regulatory network linking perturbed ligands to spatially responsive transcription factors, SCENIC+-inferred regulatory regions and predicted target genes. Ligand–TF associations indicate regulatory activity enriched near or farther from the corresponding KD cells, with edge width representing FDR q-value.

These spatial effects were also evident at individual regulatory loci. Following PDGFC KD, accessibility at representative regions increased with distance from perturbed cells, consistent with the corresponding ATAC fragment profiles (**Fig. 3f,g**). SEMA6D KD similarly produced distance-dependent changes in both accessibility and transcription-factor motif activity (**Fig. 3h**). Together, these results demonstrate coordinated cell non-autonomous transcriptional and epigenomic changes in the microenvironment of cells with perturbations targeting signaling pathways.

Finally, our dataset provides a unique opportunity to systematically probe the role that cis-regulatory elements and transcription factors play in signal transduction pathways. To investigate these questions, we used SCENIC+^20^ to connect ligand perturbations to downstream transcription factors, regulatory regions, and target genes. This generated networks linking spatially responsive transcription factors to predicted enhancer–gene relationships and their upstream ligand perturbations (**Fig. 3i**), providing a multilayer view of how local signaling perturbations propagate through downstream gene-regulatory programs.

### Cell differentiation shifts near cells with ligand perturbations

We next examined whether the distance-dependent changes in gene expression and chromatin accessibility that we observed were accompanied by changes in lineage specification and differentiation. Indeed, we did observe cell lineage specification and differentiation change as a result of perturbations. For example, WNT5A and EFNA3 KDs showed enrichment of Pluripotent and depletion of Early Ectoderm as receiver cells moved further away from these perturbations state (**Fig. 4a-c**).

**Figure 4.**
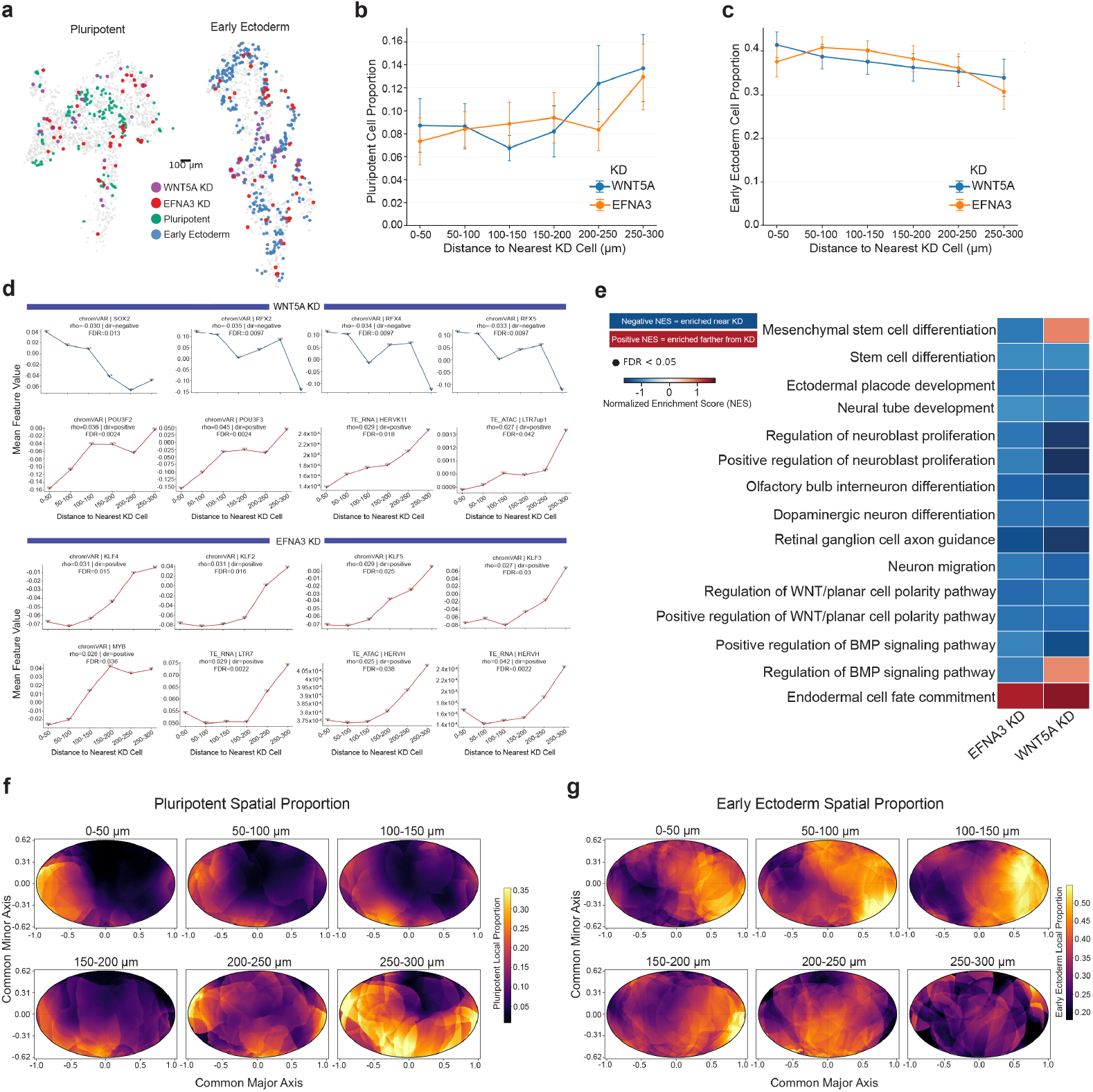
| Spatially localized ligand perturbations are associated with shifts in neighboring cell differentiation states. **a**, Spatial positions of cells in representative embryoid bodies (EBs), showing WNT5A- and EFNA3-KD cells together with Pluripotent or Early Ectoderm cells. Scale bar, 100 μm. **b**,**c**, Proportion of Pluripotent (**b**) and Early Ectoderm (**c**) receiver cells as a function of distance from the nearest WNT5A- or EFNA3-KD cell. Pluripotent-cell proportions increase with distance, whereas Early Ectoderm proportions decrease. **d**, Representative distance-dependent regulatory features following WNT5A and EFNA3 KD. WNT5A KD shows proximal enrichment of SOX2- and RFX-family chromVAR motif activity and increased pluripotency-associated POU3F and transposable-element programs with distance. EFNA3 KD shows increasing KLF-family chromVAR motif activity and HERVH/LTR7-associated transposable-element activity with distance. **e**, Targeted pre-ranked GSEA of distance-dependent transcriptional responses following EFNA3 and WNT5A KD. Genes were ranked by the magnitude and direction of their Gaussian-process response; negative normalized enrichment scores (NES) indicate enrichment near KD cells and positive NES indicate enrichment farther from KD cells. Black circles denote FDR < 0.05. **f**,**g**, Common-coordinate maps showing local Pluripotent (**f**) and Early Ectoderm (**g**) cell proportions across successive distance bins from WNT5A and EFNA3 KD cells. EBs were registered using reproducible spatial transcriptional gradients along common major and minor axes, and local cell-type proportions were estimated within individual EBs by Gaussian-kernel smoothing before averaging across EBs and perturbations. Pluripotent-cell proportions increase with distance from KD cells, whereas Early Ectoderm proportions show the reciprocal spatial pattern.

These cell-state gradients were supported by corresponding chromatin accessibility and gene expression changes. Near WNT5A KD cells, chromVAR accessibility was elevated at SOX2 and RFX2/4/5 motifs and declined with distance^21^, whereas pluripotency-associated POU3F and HERVK11/LTR7 programs increased farther from the KD^21–23^ (**Fig. 4d**). EFNA3 KD showed a similar transition, with KLF2/3/4/5 motif accessibility and HERVH/LTR7-associated transposable-element chromatin accessibility increasing with distance (**Fig. 4d**). SCENIC+ further supported these gradients, identifying increased SOX5 regulatory activity near EFNA3 KD, consistent with early ectodermal patterning^24^, and elevated NR2F1 activity near WNT5A KD, including regulation of ERBB3, while POU5F1/OCT4 activity increased with distance from WNT5A KD, consistent with greater pluripotency-associated regulatory activity farther from the perturbation^25,26^ (**Fig. 3i**). Targeted GSEA further showed enrichment of ectodermal, neural-tube and neurodevelopmental programs near both KDs, consistent with a more differentiated state proximal to the perturbations (**Fig. 4e**).

To visualize these effects across EBs, we aligned embryoid bodies using a shared spatial coordinate system based on the major and minor axis of each EB, orienting each axis using spatial gene expression gradients (**Methods**). We then mapped the locations of cells with ligand knockdowns, as well as the cell-type proportions near these knockdown cells. In this shared space, pluripotent cell proportions increased with distance from WNT5A and EFNA3 KD cells, whereas Early Ectoderm proportions decreased (**Fig. 4f,g**). Interestingly, pluripotent cells were preferentially localized toward the negative end of the minor axis, enriched for embryonic stem-cell and primitive epithelial programs. Early Ectoderm cells were preferentially localized toward the positive end of the major axis, associated with WNT-related transcriptional programs. Together, these results demonstrate shifts in the differentiation states in the spatial neighborhood of cells with ligand knockdowns.

### Kinase perturbation effects vary with local ligand-receptor signaling

Having interrogated extracellular ligands and membrane-bound receptors, we next used Perturb-Tags to study the effects of perturbing kinases–the intracellular layer of signal transduction pathways. We constructed a pooled CRISPRi library containing five sgRNAs targeting all of the approximately 300 kinases expressed in the EB system. To achieve sufficient representation across this large library, we profiled nine Trekker slides across thirteen 10X Chromium lanes, yielding 70,235 cells across 806 EBs **(Fig. 5a)**. EBs in the kinase screen were generally smaller than those in the ligand and receptor experiments, consistent with broad effects of kinase perturbation on growth, survival and differentiation **(Fig. 5b)**. Nevertheless, guides targeting most kinases were recovered, and several perturbations formed locally expanded guide-positive patches **(Extended Data Fig. 7)**. Kinases associated with large guide-positive clones included TTBK2, PTK2/FAK, MAPK8/JNK1, ROCK2, PNCK, and JAK3, several of which have established roles in regulating cell proliferation, survival, and apoptosis^27–33^, suggesting that their knockdown may confer relative growth or survival advantages during EB development.

**Figure 5:**
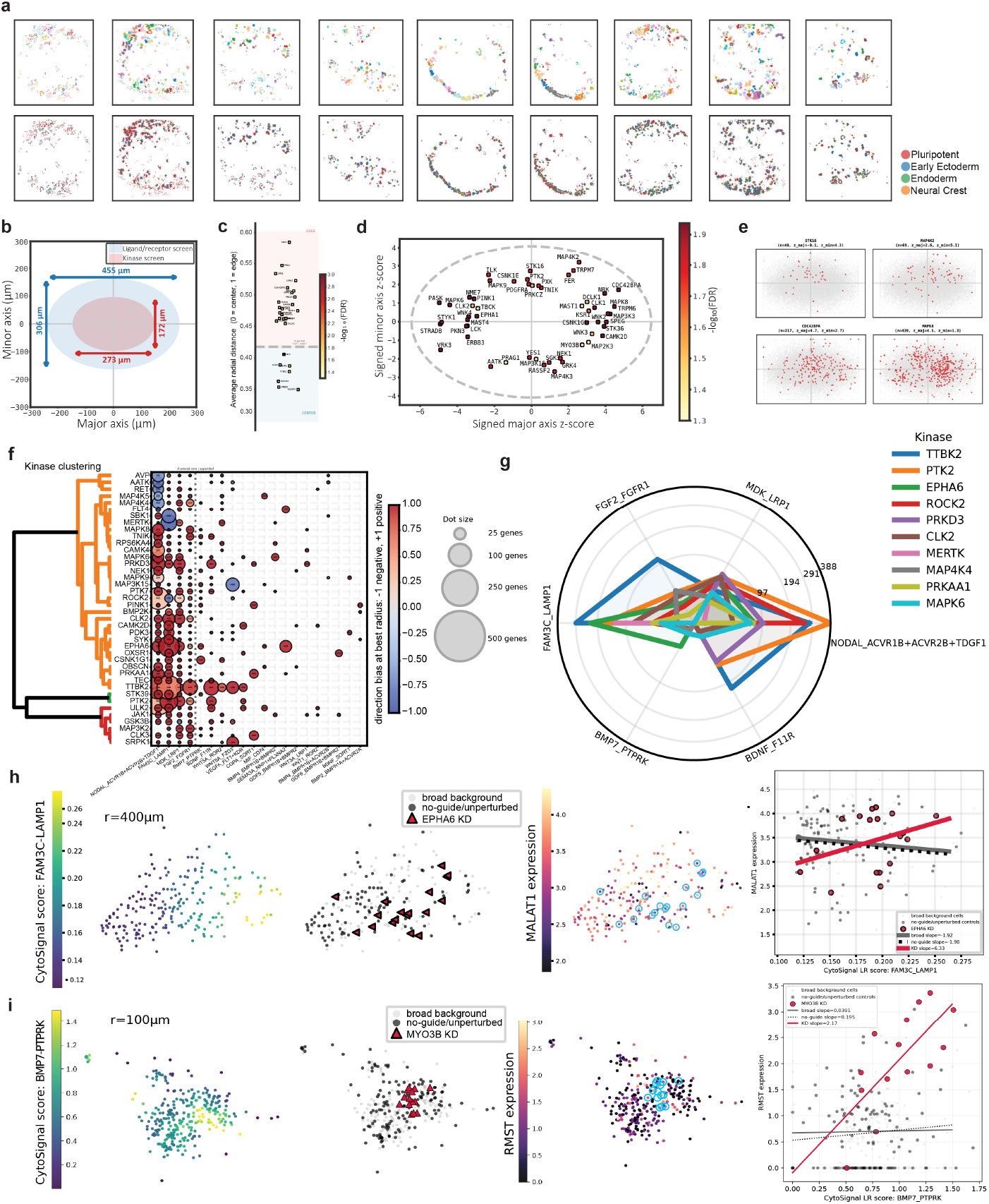
Perturb-Tags reveals context-specific kinase perturbation effects. **a**, Spatial plot of cell locations colored by cell type (**top**) and embryoid body assignment (**bottom**) from 9 spatial transcriptomic slides. **b**, Schematic comparing the spatial footprints sampled in the ligand/receptor screen (blue; approximately 455 × 306 µm) and kinase screen (red; approximately 273 × 172 µm). **c**, Average radial position of cells carrying individual kinase perturbations relative to the EB center. The dashed line indicates the screen-wide reference, with perturbations above or below the reference showing preferential localization toward the EB edge or center, respectively; color indicates −log10(FDR). **d**, Spatial distribution of kinase perturbations along the EB major and minor axes, represented as signed z-scores. Points are colored by −log10(FDR), and the dashed ellipse indicates average dimension of the fitted ellipses for all EBs. **e**, Representative spatial distributions of STK16-, MAP4K2-, CDC42BPA-, and MAPK8-perturbed cells, with kinase-KD cells shown in red. **f**, Clustered bubble plot summarizing significant kinase-KD × incoming ligand–receptor (LR) interaction effects obtained from linear regression models of gene expression as a function of kinase knockdown, local CytoSignal LR score, and their interaction. Rows represent kinase perturbations and columns represent LR scores. Bubble size indicates the number of genes with significant kinase-KD × LR interactions, while color indicates the overall direction of the interaction profile, from predominantly negative (blue) to predominantly positive (red). Hierarchical clustering groups kinases with similar signed interaction profiles across LR contexts. **g**, Radar plot showing the numbers of significant interaction genes for the ten strongest kinase candidates across six prominent LR contexts: NODAL– ACVR1B/ACVR2B/TDGF1, FAM3C–LAMP1, MDK–LRP1, FGF2–FGFR1, BMP7–PTPRK, and BDNF–F11R. Distinct kinase-specific profiles demonstrate that sensitivity to local LR signaling varies across perturbations. **h**, Representative MAP3K15-KD example. From left to right: spatial distribution of the incoming NODAL_ACVR1B+ACVR2B+TDGF1 CytoSignal score calculated at a 100-µm radius; locations of MAP3K15-KD cells relative to broad-background and no-guide/unperturbed cells; spatial expression of RYR2 with MAP3K15-KD cells highlighted; and interaction plot relating NODAL_ACVR1B+ACVR2B+TDGF1 score to RYR2 expression. RYR2 expression increases with NODAL_ACVR1B+ACVR2B+TDGF1 signaling in MAP3K15-KD cells, whereas the relationship is less positive in surrounding background cells. **i**, Representative MYO3B-KD example shown analogously for the incoming BMP7– PTPRK CytoSignal score at a 100-µm radius and RMST expression. RMST expression increases strongly with BMP7–PTPRK signaling in MYO3B-KD cells, whereas the corresponding relationship is weak or nearly flat in background cells. In **h–i**, red lines indicate kinase-KD-specific regression fits, gray lines indicate broad-background fits, and black dotted lines indicate no-guide/unperturbed-control fits.

We also identified notable patterns in the spatial distribution of cells with kinase knockdowns. As with the Multiome data, we mapped the nuclei recovered from the kinase dataset into a spatial coordinate system based on oriented major and minor axes for each embryoid body (**Methods**). Despite the smaller size of the EBs in the kinase dataset, we again noticed that pluripotent and early ectoderm cells tended to occupy opposing ends of the major axis. Using a spatial permutation approach, we identified kinases that tended to be located more centrally or peripherally in EBs than expected by chance (**Fig. 5c**). We then performed a more detailed spatial permutation analysis to identify kinase perturbations with spatial enrichment in either the positive or negative direction of the major or minor axis (**Fig. 5d-e**). These analyses indicated that cells with certain kinase knockdowns indeed occupied preferential positions along the major and minor axes, suggesting that the kinases involved may influence spatial patterning of the embryoid body.

We next tested whether kinase perturbation effects depended on local upstream signaling. Gene expression was modeled as a function of kinase knockdown, spatial CytoSignal LR scores, and their interaction. Significant kinase-KD × LR-score interactions therefore identify transcriptional effects that vary with the local signaling environment. Across the screen, kinases differed in the number, direction, and spatial scale of significant interactions, with clustering revealing distinct context-dependent response profiles **(Fig. 5f)**.

Among six prominent LR interactions, TTBK2 and PTK2 showed the largest numbers of significant interaction genes, followed by EPHA6, ROCK2, PRKD3, CLK2, MERTK, MAP4K4, PRKAA1 and MAPK6 **(Fig. 5g)**. NODAL- and FAM3C-associated signals produced particularly strong interactions, whereas MDK, FGF2, BMP7 and BDNF showed more selective kinase-specific effects. Visual inspection confirms these context-specific perturbation effects. For example, RYR2 expression was more strongly associated with the 100-µm incoming NODAL– ACVR1B/ACVR2B/TDGF1 score in MAP3K15-KD cells than in surrounding background cells **(Fig. 5h)**. Similarly, RMST increased with the 100-µm incoming BMP7–PTPRK score in MYO3B-KD cells, whereas this relationship was nearly absent in surrounding cells **(Fig. 5i)**. Together, these results suggest that factors in the microenvironment of cells modulate the transcriptional consequences of kinase perturbation.

### Graph attention networks predict tissue-wide effects of genetic perturbation on gene expression and chromatin accessibility

Having observed spatially distributed cell-non-autonomous effects across gene expression, LR signaling, chromatin state, and differentiation, we next tested whether these responses were sufficiently structured, reproducible, and generalizable to be learned across independent tissues. We developed a graph neural network model that predicts the effects of perturbations on gene expression and chromatin accessibility across an entire tissue. The model takes a graph as input. Each nucleus is represented as a node and spatially proximal nuclei within the same EB are connected by edges. The model takes spatial positions, CRISPR guide assignments, and cell-type identities as inputs and predicts gene expression and chromatin accessibility of each nucleus as outputs. A non-spatial multilayer perceptron (MLP) first learns the cell-intrinsic effects of each guide RNA and cell type. The graph attention network learns a spatial perturbation correction from neighboring cells, which is then combined with the cell-intrinsic prediction. Separate RNA and ATAC branches predict gene expression and chromatin accessibility features (**Fig. 6a**).

**Figure 6:**
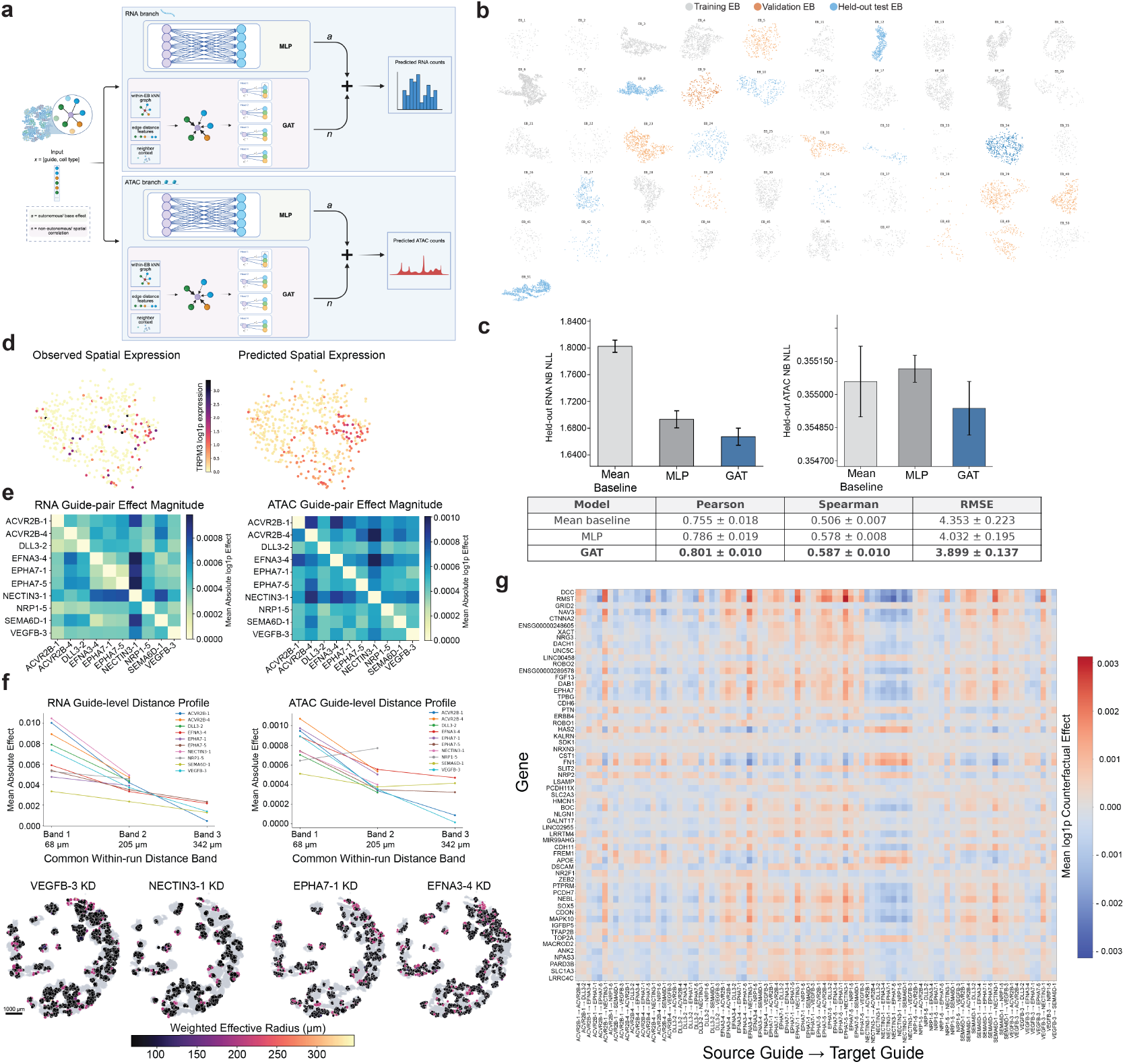
Graph attention network model predicts cell-non-autonomous gene expression and chromatin accessibility responses. **a**, Architecture of the modality-specific spatial-residual graph attention model. For RNA and ATAC, a non-spatial multilayer perceptron (MLP) predicts counts from each nucleus’s guide assignments and cell-type identity, while a distance-aware graph attention network (GAT) learns a non-autonomous spatial correction from neighboring guide and cell-type identities. Final predictions combine the autonomous base effect (*a*) and non-autonomous spatial correction (*n*). **b**, Spatial distribution of nuclei across EB-level training (grey), validation (orange) and held-out test (blue) assignments used across ten held-out-EB evaluations. **c**, Held-out RNA and ATAC negative-binomial negative log-likelihood (NB NLL) for the MLP and GAT. Error bars indicate paired s.e.m. across held-out EBs. Table shows held-out RNA prediction performance for the mean baseline, MLP and GAT; values are mean ± s.e.m. across EBs. **d**, Observed and GAT-predicted spatial pattern of TRPM3 log1p expression in a representative held-out EB. **e**, RNA and ATAC counterfactual effect magnitudes following guide-to-guide substitution in neighboring cells. Rows denote the source guide and columns the target guide; color indicates the mean absolute log1p change in the predicted receiver-cell state. **f**, Distance-resolved counterfactual effects obtained by replacing the perturbation guide in neighboring guide-positive cells with the non-targeting control guide separately within successive spatial distance bands, while leaving the receiver cell unchanged. Top, mean absolute RNA and ATAC effects for individual perturbation guides across common within-run distance bands. Bottom, representative spatial maps of perturbation-specific effective radii. Colored circles denote receiver cells with modeled non-autonomous effects; color indicates the effect-weighted effective radius (**Methods**) and circle size indicates effect magnitude. Scale bar, 1,000 µm. **g**, Gene-level RNA responses to ordered source-to-target guide substitutions in neighboring cells. Color indicates the mean log1p counterfactual change in predicted receiver-cell expression, with positive values indicating increased and negative values decreased predicted expression following guide substitution. RNA genes were selected on the basis of biological variability rather than counterfactual effect magnitude.

We evaluated the model across 10 held-out EBs, using separate validation EBs for model selection (**Fig. 6b**). The addition of spatial information improved prediction of gene expression relative to the non-spatial MLP, with a pearson correlation of 0.652 in predicting RNA counts (**Extended Data Fig. 8c**). Similarly, the spatial model achieved a better ATAC loss compared to the non-spatial model (**Fig. 6c**). Predicted and observed guide-averaged expression profiles were also highly correlated for predicting both ATAC and RNA (r = 0.801). Note that all of these metrics were calculated across the held-out EB predictions rather than the training data. The learned graph therefore captured predictive information from neighboring source cells beyond the guide and cell-type information of the target cell alone. Visual inspection confirms that the GAT reproduces the spatial patterns of individual genes in held-out EBs. (**Fig. 6d**).

We then used the trained GAT for counterfactual inference to determine how perturbation effects propagate through spatial neighborhoods. We first tested how changing the identity of a neighboring perturbation altered the predicted state of receiver cells. Neighboring cells carrying one guide were changed *in silico* to a second guide, while receiver cells carrying either guide were excluded and remained unmodified. Different guide substitutions produced distinct RNA and ATAC effect magnitudes, indicating that neighboring perturbation identity shapes the predicted non-autonomous molecular response (**Fig. 6e**).

We next examined how far these non-autonomous effects extended through space. For each perturbation, neighboring guide-positive cells were grouped into distinct spatial distance bands, and within each band the perturbation guide was replaced *in silico* with the non-targeting control guide while the receiver cell remained unchanged; the resulting changes in predicted RNA and ATAC states were then quantified separately for each distance band. For most perturbations, the predicted effect was strongest in proximal neighborhoods and decreased with distance in both RNA and ATAC (**Fig. 6f**). The spatial maps showed that the modeled non-autonomous effects were heterogeneous across the tissue, with different receiver cells exhibiting both different effect strengths and different effective spatial ranges (**Fig. 6f, Extended Data Fig. 8d**). For example, VEGFB KD showed predominantly short-range RNA effects across receiver cells, with scattered cells exhibiting intermediate effective radii of roughly 150– 250 µm, whereas EFNA3 KD showed a similarly heterogeneous spatial distribution of ATAC effects.

At the gene level, guide-to-guide counterfactual substitutions revealed how changing the identity of neighboring perturbations altered receiver-cell expression. Distinct source-to-target guide swaps produced gene-specific increases and decreases in predicted expression (**Fig. 6g**). Together, these results show that cell-non-autonomous perturbation effects are sufficiently structured and reproducible for spatial context to improve prediction across held-out tissues and to model how perturbation identity and distance shape neighboring-cell responses.

## Discussion

Understanding multicellular development requires resolving not only the cell-intrinsic consequences of genetic variation, but also how perturbations in one population alter the molecular state and developmental trajectory of surrounding cells. Pooled CRISPR and Perturb-seq approaches have enabled systematic interrogation of cell-intrinsic gene function, but dissociation removes tissue architecture and therefore obscures the spatial relationships through which many intercellular signals operate^7,34^. Spatial functional-genomics approaches have begun to address this limitation by preserving the relationship between perturbation identity and tissue context^10,12,35^. Here, we developed Perturb-Tags, a spatial multi-omic perturbation platform that links CRISPR guide identity with gene expression, chromatin accessibility and spatial position. Applying Perturb-Tags to human embryoid bodies across LR and kinase perturbation screens revealed widespread distance-dependent cell-non-autonomous responses and connected these responses to transcriptional regulation, chromatin state, cell differentiation and the spatial organization of developing tissue. These findings establish a framework for measuring how localized genetic perturbations propagate through neighboring cell populations rather than considering the perturbed cell as an isolated unit.

Several features of the screens provided direct evidence that Perturb-Tags captures such spatially distributed responses. Specific perturbations formed expanded spatial clones, consistent with the principle that perturbations affecting cellular fitness can become enriched during pooled CRISPR selection^35^. Importantly, cells surrounding these clones exhibited transcriptional changes as a function of their distance from cells carrying the perturbation even when the receiver cells did not themselves carry the perturbation being analyzed. LR perturbations provided a particularly clear example: direct LR interactions involving the targeted signaling component were reduced proximal to the corresponding KD and recovered with increasing distance, whereas a distinct set of compensatory LR interactions showed the opposite spatial pattern. The same organization extended beyond transcriptional signaling scores. Independent Multiome measurements identified distance-dependent changes in chromatin accessibility, gene activity and transcription-factor motif activity around ligand perturbations, indicating that localized perturbations can generate coordinated transcriptional and epigenomic responses in surrounding cells.

To extend these observations beyond individual perturbation–response relationships, we developed a spatial graph attention network (GAT) in which each nucleus was modeled together with its local cellular neighborhood. GATs provide a natural framework for spatial tissue data because they allow individual neighbors to contribute differently to the representation of a receiver node^36^. Our model separated the cell-intrinsic prediction from the spatial contribution of neighboring cells and improved prediction of RNA and ATAC states relative to a non-spatial MLP across held-out EBs. Counterfactual perturbation of neighboring guide identities subsequently identified guide-pair-specific changes in predicted receiver-cell states, while distance-resolved counterfactual analysis showed that many predicted perturbation effects were strongest in proximal neighborhoods and weakened with distance. Previous perturbation studies established that genetic perturbations can alter the states of other cells and recognized paracrine signaling as a major limitation of pooled screens, but generally lacked the spatial information required to determine how these effects propagate through tissue^10,37^.

More broadly, these results illustrate how spatial perturbation assays can be used to screen for complex, multicellular phenotypes in tissues rather than single-cell phenotypes in cell culture systems. Recent approaches including Perturb-map, Spatial Perturb-seq and Perturb-DBiT similarly demonstrate the value of retaining tissue architecture during pooled genetic screening^10,12,34^, but the increased sensitivity of Perturb-Tags and integration of spatial perturbation identity with transcriptomic and epigenomic measurements provide new opportunities to connect extracellular perturbations to the regulatory programs through which receiver cells change state. We anticipate that Perturb-Tags could be used to examine how genetic perturbations reshape cell–cell communication and differentiation in organoids, developmental models, and disease-associated tissue microenvironments. Extending these experiments across developmental time, additional types of perturbations, and more complex human tissue models should also help distinguish transient signaling responses from persistent changes in cell identity. Together, Perturb-Tags provides a generalizable framework for determining how localized genetic perturbations propagate through multicellular environments to reshape the molecular and developmental states of neighboring cells.

**Extended Data Figure 1:**
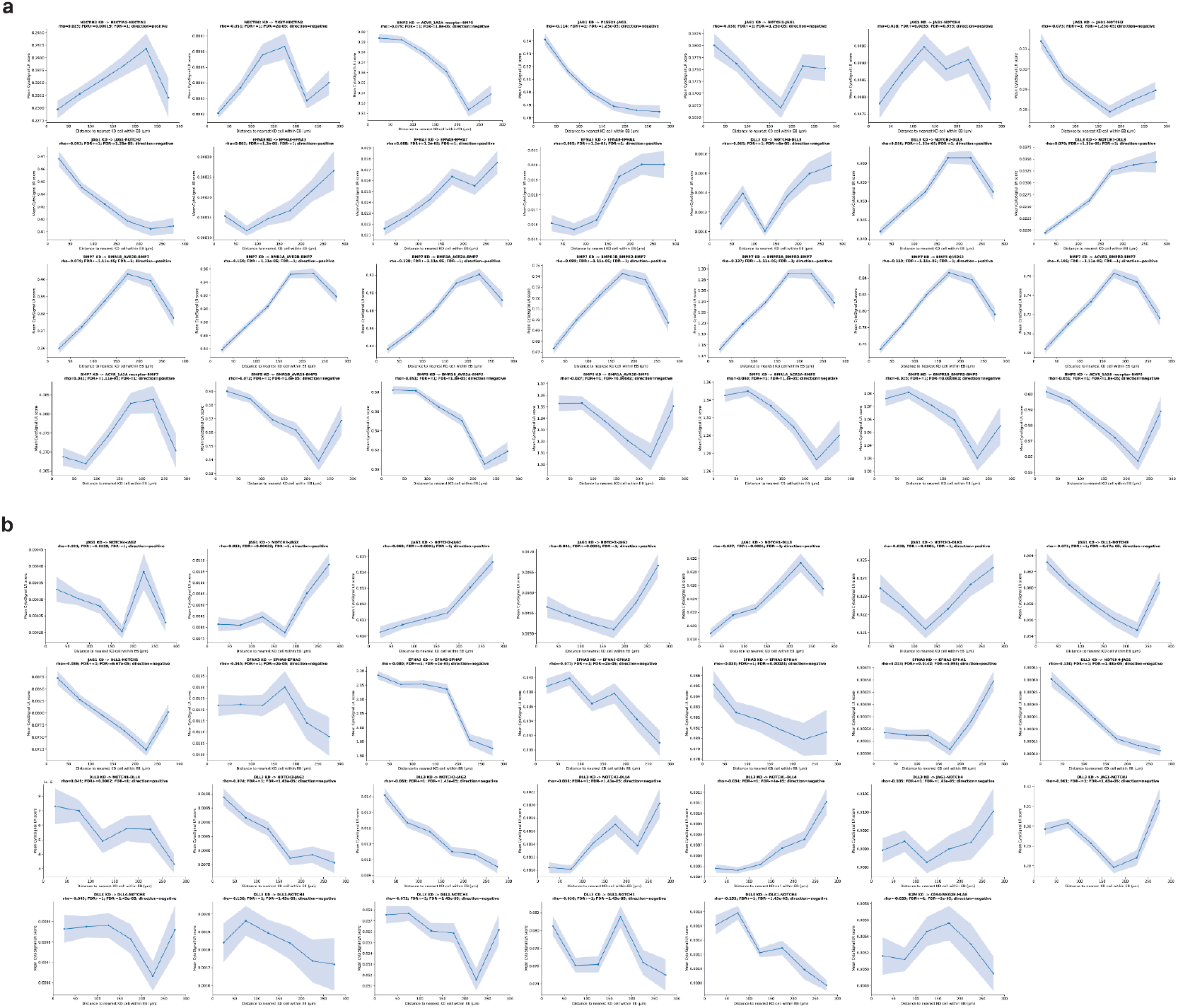
Distance-dependent direct and compensatory ligand–receptor signaling responses following CRISPRi perturbation. **a**, CytoSignal LR scores for direct interactions plotted as a function of distance from the nearest cell carrying the corresponding perturbation within each EB. Direct interactions contain the targeted signaling component itself. **b**, Distance-dependent CytoSignal LR scores for compensatory interactions involving related or alternative signaling components following perturbation. Lines and points indicate mean LR score and shaded regions indicate s.e.m. Spearman’s q and direction-specific FDR values for positive (FDR+) and negative (FDR−) distance associations are indicated above each plot.

**Extended Data Figure 2:**
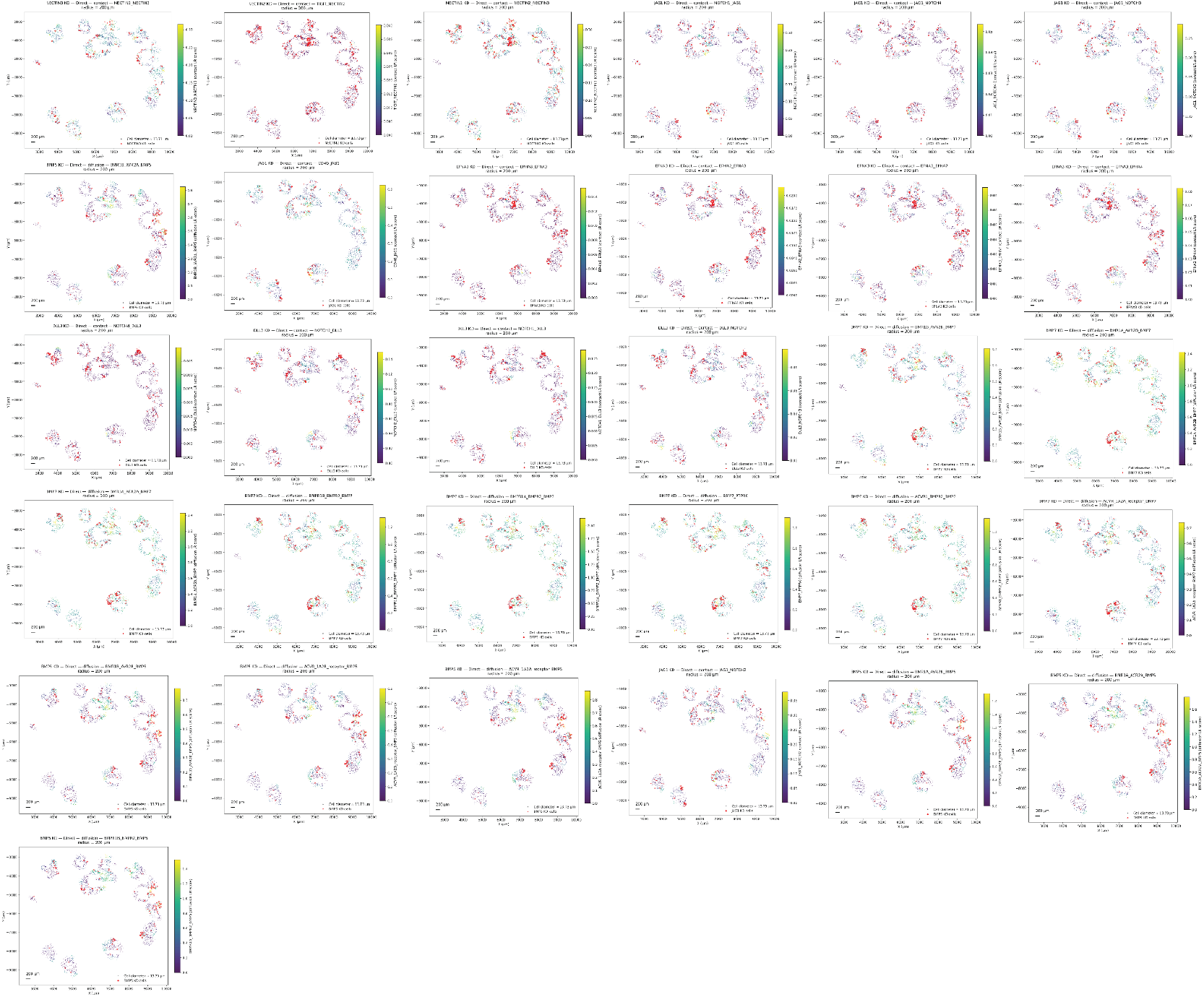
Spatial distribution of direct ligand–receptor signaling responses around perturbed cells. Spatial maps of representative direct CytoSignal LR interactions identified following CRISPRi perturbation. Receiver cells are colored according to the CytoSignal LR score of the indicated contact- or diffusion-mediated interaction, and red points indicate cells carrying the corresponding perturbation. Titles indicate the perturbation, interaction class and LR pair. Maps were evaluated within a 200-µm spatial radius. Scale bars, 200 µm.

**Extended Data Figure 3:**
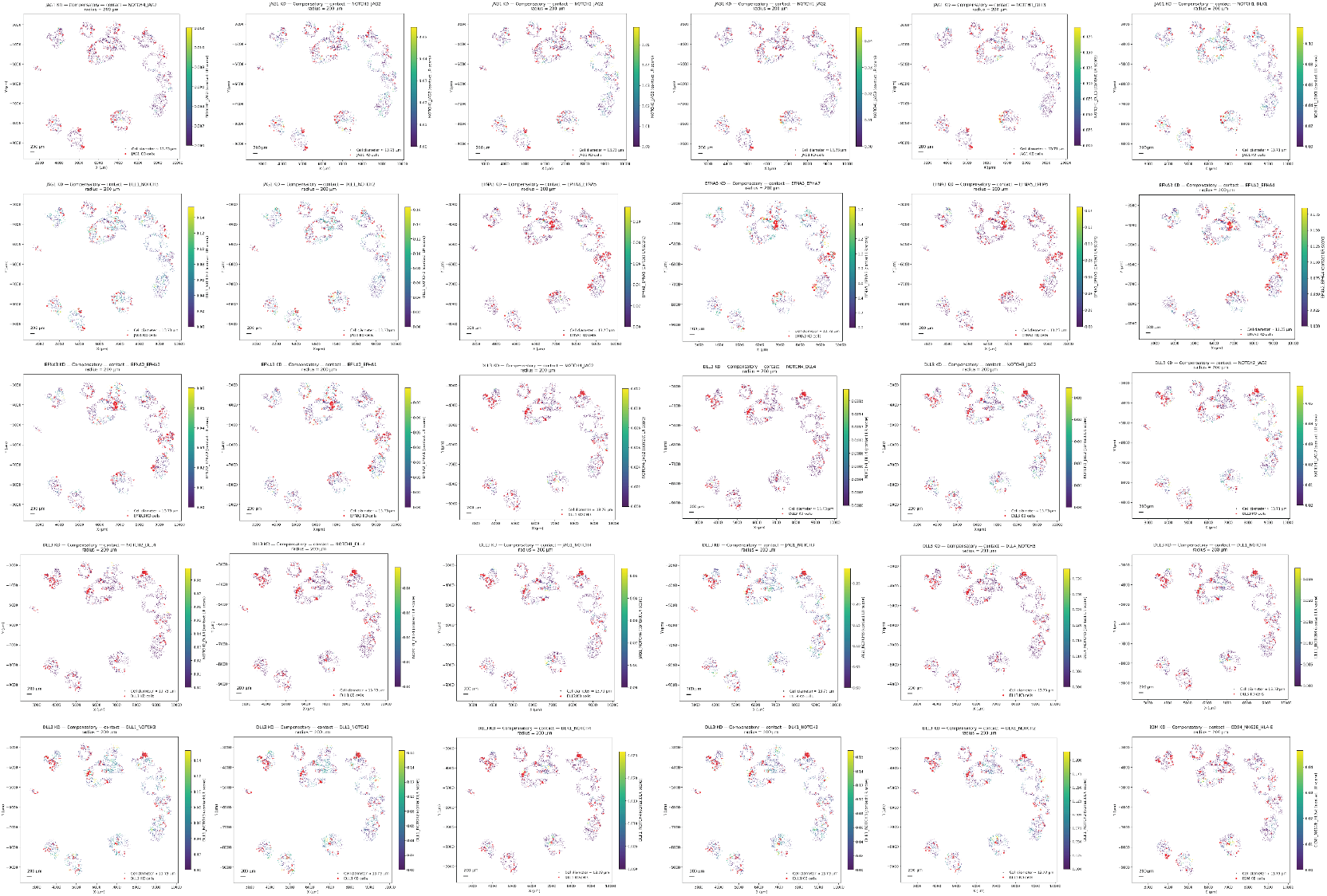
Spatial distribution of compensatory ligand–receptor signaling responses around perturbed cells. Spatial maps of compensatory CytoSignal LR interactions following CRISPRi perturbation, including alternative ligand–receptor interactions within related signaling pathways. Receiver cells are colored according to the CytoSignal LR score of the indicated interaction, and red points indicate cells carrying the corresponding perturbation. Titles indicate the perturbation, interaction class and LR pair. Maps were evaluated within a 200-µm spatial radius. Scale bars, 200 µm.

**Extended Data Figure 4:**
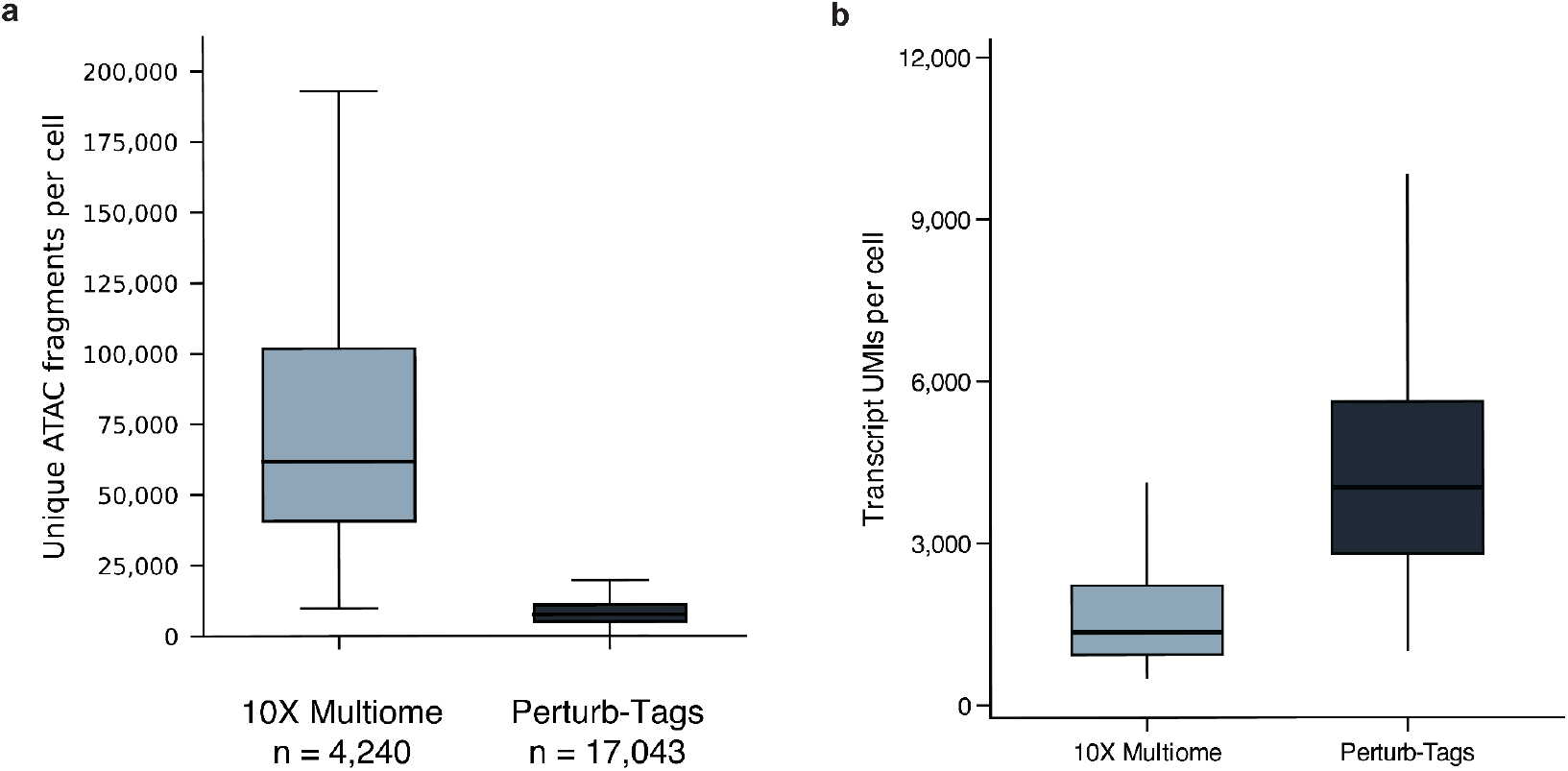
Comparison of per-cell ATAC and transcript capture between 10X Multiome and Perturb-Tags. **a**, Unique ATAC fragments per cell for 10X Multiome (n = 4,240 cells) and Perturb-Tags (n = 17,043 cells). **b**, Transcript UMIs per cell for the same cell cohorts. Centre lines indicate medians; box limits indicate the 25th and 75th percentiles; whiskers extend to 1.5 times the interquartile range. Outliers are not shown.

**Extended Data Figure 5:**
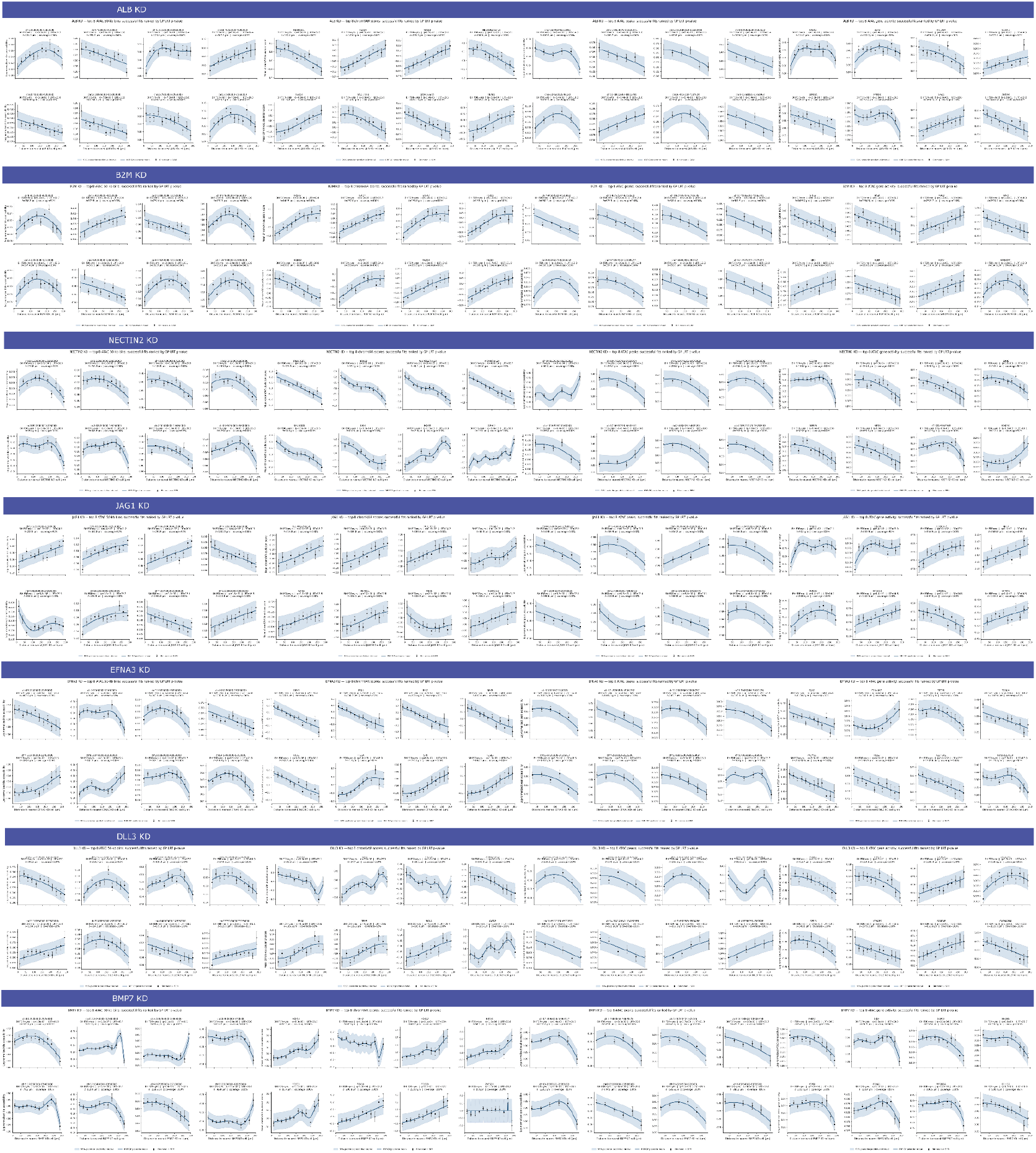
Gaussian-process modeling identifies distance-dependent chromatin responses to perturbation. Distance-dependent regulatory responses following ALB, B2M, NECTIN2, JAG1, EFNA3, DLL3 and BMP7 KD across four ATAC-derived modalities: fixed 50-kb ATAC bins, chromVAR motif-deviation scores, ATAC peaks and ATAC gene-activity scores. For each perturbation and modality, the eight successfully fitted features with the strongest Gaussian-process (GP) likelihood-ratio-test evidence are shown. Black points and error bars indicate distance-bin means ± s.e.m.; dark-blue curves indicate the radial-basis-function GP posterior mean and shaded regions indicate the 95% posterior predictive interval. Subplot annotations report Benjamini–Hochberg FDR status and q value, likelihood-ratio statistic, fitted GP length scale (*l*) and predictive coverage.

**Extended Data Figure 6:**
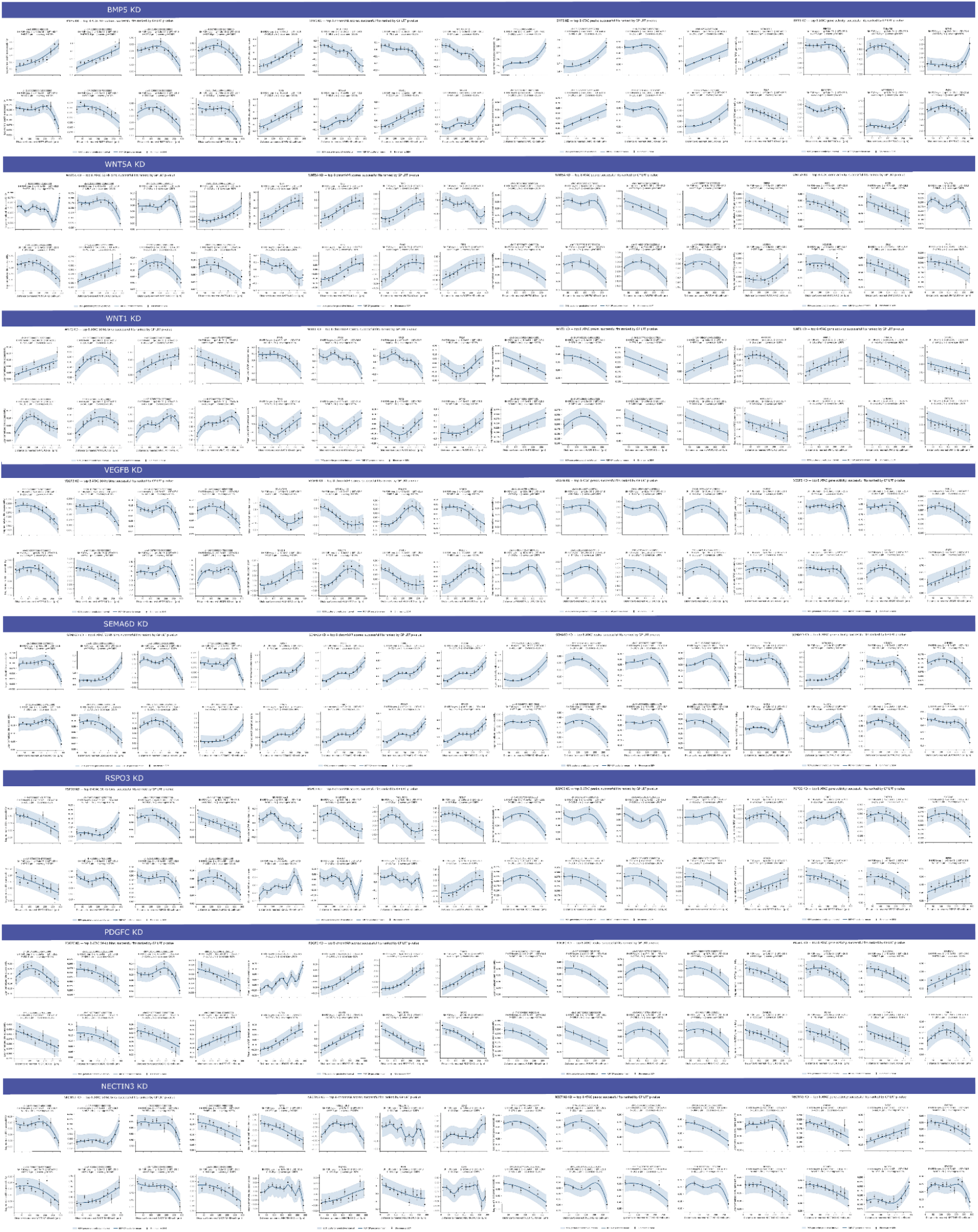
Additional Gaussian-process models of distance-dependent chromatin responses to perturbation. Distance-dependent regulatory responses following BMP5, WNT5A, WNT11, VEGFB, SEMA6D, RSPO3, PDGFC and NECTIN3 KD across fixed 50-kb ATAC bins, chromVAR motif-deviation scores, called ATAC peaks and ATAC gene-activity scores.

**Extended Data Figure 7:**
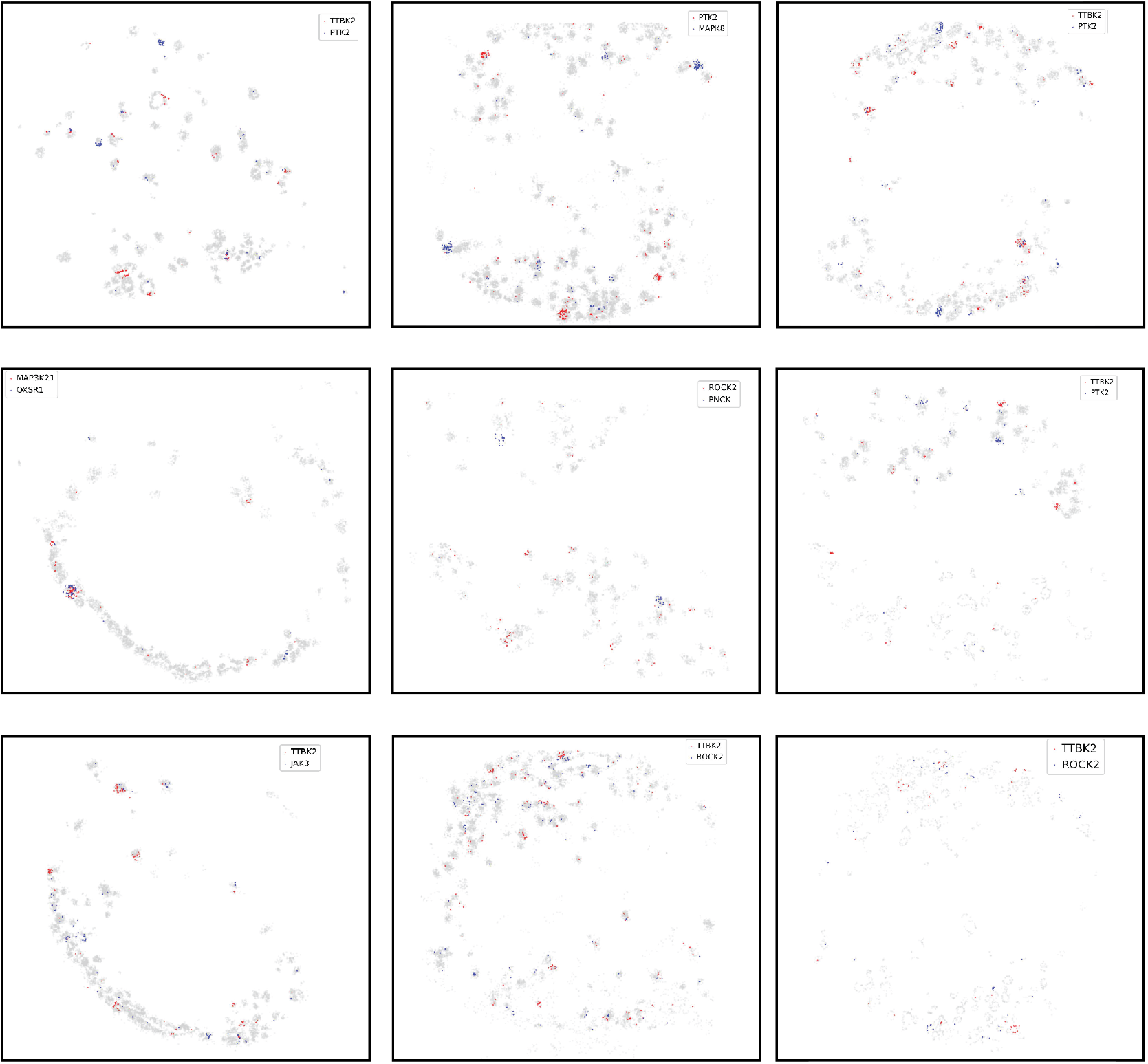
Representative spatial distributions of clonally expanded perturbations. Spatial maps showing the distribution of pairs of Kinase CRISPRi perturbations across representative EBs. Red and blue points indicate nuclei carrying the two indicated perturbations in each map, whereas grey points indicate the remaining nuclei. The maps illustrate the spatially localized and heterogeneous organization of independently expanded perturbation populations.

**Extended Data Figure 8:**
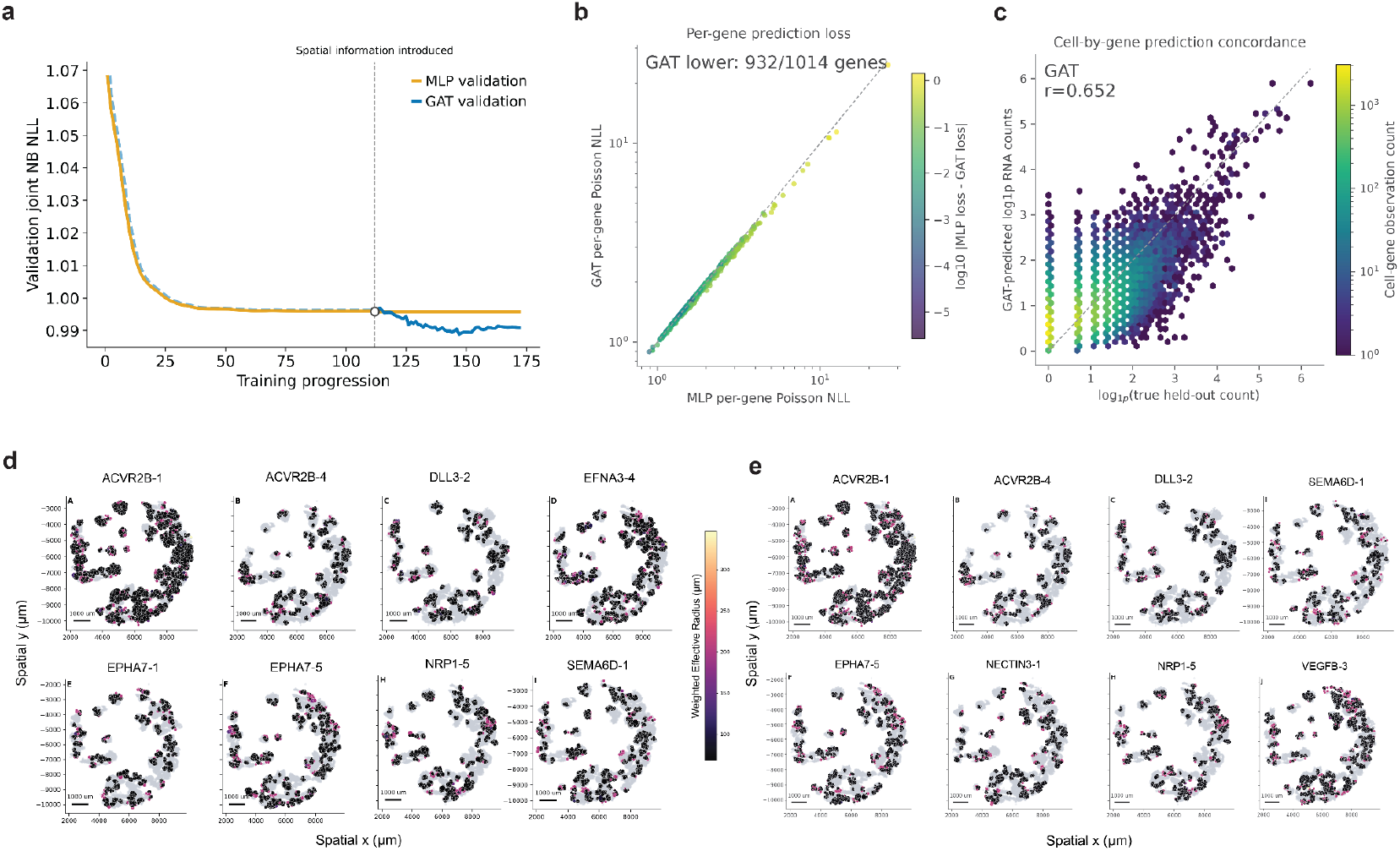
Validation and spatial characterization of the graph-attention model. **a**, Representative validation joint negative-binomial negative log-likelihood (NB NLL) during non-spatial MLP training and subsequent spatial-residual GAT training. The dashed vertical line indicates the point at which spatial information was introduced and GAT spatial parameters were trained. **b**, Per-gene held-out RNA prediction loss for the MLP and GAT. Dashed line indicates equal performance; the GAT has lower Poisson NLL for 932 of 1,014 evaluated genes, and color indicates the log-transformed magnitude of the difference between MLP and GAT loss. **c**, Cell-by-gene concordance between observed held-out RNA counts and GAT predictions on the log1p scale. Dashed line indicates equality; Pearson’s *r* = 0.652 and color indicates the number of cell–gene observations. **d**,**e**, Spatial maps of GAT-inferred effect-weighted effective radii for representative perturbations in RNA (**d**) and ATAC (**e**) across pooled EBs. Grey points indicate background nuclei and colored points indicate receiver cells with modeled non-autonomous responses; color indicates the effect-weighted effective radius and point size indicates the magnitude of the predicted effect. RNA effective radii were calculated using variance-selected informative genes, whereas ATAC radii were calculated from the modeled ATAC features. Scale bars, 1,000 µm.

## Methods

### iPSC maintenance

iPSCs stably expressing GCaMP6f were cultured in mTeSR1 (STEMCELL Technologies, 85850) on dishes with Matrigel (Corning, 354230). When iPSCs reached 70-80% confluency, cells were passaged by washing with DPBS (Gibco, 14190144), and dissociated with Accutase (Gibco, A1110501) at 37°C for 5 minutes. Accutase was then diluted 1:2 in DPBS, and the cells were collected in conical tubes and centrifuged at 300xg for 3 minutes. The supernatant was aspirated and resuspended with mTeSR1 supplemented with ROCK inhibitor (Stemcell, NC1678100). The cells were counted and plated onto Matrigel-coated plates at the appropriate number.

### Molecular cloning and gRNA library

Lentiguide(10x)-BFP-Puro was generously gifted by Dr. Gary Hon from UT Southwestern Medical Center. We chose to perturb 25 ligands and receptors showing high expression in our previously published Embryoid Bodies (EB) RNA velocity data [topovelo] as well as 301 kinases that are highly differentially expressed in the same dataset, while excluding cell cycle genes. We selected 5 previously validated guide RNAs44 per target gene, plus 5 additional non-targeting guides (Negative-Controls). The oligonucleotide pool (Twist Bioscience) was PCR amplified and cloned using Gibson assembly into the single gRNA expression plasmid. The transformed plasmids were electroporated into Endura DUOs electrocompetent cells (LGC Biosearch Technologies, 60242-1). Transformed cells were added to LB broth supplemented with Ampicillin and incubated for 12-16 hours at 37°C in a shaker, and then proceeded to Maxi prep for plasmid DNA extraction (QIAGEN, 12162). Whole plasmid sequencing (Plasmidsaurus) was performed to verify that gRNAs were cloned successfully.

### HEK293T culture and lentivirus production

HEK293T cells were cultured in DMEM (Gibco, 11995065) supplemented with 10% FBS. Once the cells reached 80% confluency, they were dissociated using TrypLE Express (Gibco, 12604013) for 3 minutes at 37°C. TrypLE was diluted with DPBS and centrifuged for 3 minutes at 300g. Cells were counted and seeded onto a plate at the appropriate number. For lentivirus production, 10-cm dishes were seeded with 7 x 106 HEK293T cells in 12 mL of lentivirus packaging medium (Opti-MEM I (Gibco, 31985070), 5% FBS, 200 μM sodium pyruvate (Gibco, 11360070)). The next day, the transfection was performed using the following protocol by combining transfection mixes A and B. Transfection mix A contained 10 μg CRISPRi library and 26 μL of Lentiviral Packaging Mix (MISSION, SHP001) was mixed with 1.5 ml of available under a CC-BY-NC-ND 4.0 International license. (which was not certified by peer review) is the author/funder, who has granted bioRxiv a license to display the preprint in perpetuity. It is made bioRxiv preprint doi: https://doi.org/10.1101/2025.11.03.686351; this version posted November 5, 2025. The copyright holder for this preprint Opti-MEM I and 36 ul of P3000 enhancer reagent (Invitrogen, L3000015). Transfection mix B contained 1.5 ml of Opti-MEM I and 42 μL of Lipofectamine 3000 reagent (Invitrogen, L3000015). Transfection mixes A and B were combined and incubated for 20 minutes at room temperature. During incubation, 6 mL of lentivirus packaging medium was removed from the 10 cm plate. The transfection mixes A and B were gently added and mixed. Six hours later, the Lipofectamine-containing medium was carefully removed and replaced with fresh lentivirus packaging medium. Lentivirus-containing medium was collected after 24 hours and stored at 4 °C. 12 ml of fresh lentivirus packaging medium was carefully added. Lentivirus-containing medium was collected again the next day. Combined lentivirus-containing medium (approximately 24 ml) was filtered through a 0.45 μm filter to remove cell debris into a new 50 ml conical tube. After filtering, ⅓ volume of Lenti-X Concentrator (Takara, 631231) was added to the filtered solution, mixed well, and stored at 4°C overnight. Following incubation, the solution was centrifuged at 4°C for 45 minutes at 1,500g. The supernatant was removed carefully, and the virus pellet was resuspended with 1 mL mTeSR and stored at -80°C until further analysis

### EB differentiation and maintenance

For EB differentiation of CRISPRi KD iPSCs, the EB formation medium (DMEM/F-12 (Gibco, 11320033), 20% (v/v) Knockout Serum Replacement (Gibco, 10828028), 1% MEM non-essential amino acid solution (Gibco, 11140050), 0.5% GlutaMAX (Gibco, 35050061) supplemented with β-mercaptoethanol (MP Biomedicals, 194705) and 1% penicillin–streptomycin (Gibco, 15140122)) was used following the STEMCELL AggreWell 400 protocol. The AggreWell 400 24-well plate (STEMCELL Technologies, 34415) was coated with anti-adherence rinsing solution (STEMCELL Technologies, 07010). The iPSCs were dissociated with Accutase and seeded 1.2 × 10^6^cells per well in EB formation medium. After 24 hours, 50% of the medium was replaced with fresh EB formation medium. After 48 hours of seeding, the EBs were harvested using 37-µm reversible strainers and moved to an ultra-low attachment six-well plate (Corning Costar, 07-200-601) in Essential 6 (E6) medium (Gibco, A1516401). The medium change was done every 48 hours. After 14 days of seeding, the EBs were flash frozen in OCT media and kept in -80 until further analysis.

### Cryosectioning, spatial barcoding, and single-nucleus sequencing

Flash-frozen, OCT-embedded EBs were cryosectioned at 10 µm thickness for hematoxylin and eosin (H&E) staining and at 30 µm thickness for placement onto Takara Bio Trekker tiles. Following tissue placement, spatial barcoding was performed according to the Takara Bio Trekker Single-Nuclei Spatial Mapping Kit protocol (ref). Spatial barcodes were photocleaved using UV illumination, enabling their attachment to adjacent tissue regions positioned on the tile. Following spatial barcoding, nuclei were isolated using Nuclei Isolation Buffer V2 and processed using the appropriate 10x Genomics single-nucleus workflow. Library prep and next-generation sequencing were carried out in the BRCF Advanced Genomics Core at the University of Michigan. Research reported in this publication was supported by the National Cancer Institute of Health under Award Number P30CA046592 by the use of the following Cancer Center Shared Resource: Single Cell and Spatial Analysis Shared Resource. For the first two perturbation experiments, targeting ligand–receptor (LR) and kinase genes respectively, libraries were generated using the 10x Genomics Chromium Next GEM Single Cell 3’ v3.1 workflow with Feature Barcoding for CRISPR screening. Libraries from the LR perturbation experiment were sequenced on an Illumina NovaSeq X using a 10B 100 cycle flow cell, whereas libraries from the kinase perturbation experiment were sequenced on the Ultima Genomics UG 100 Single-Wafer Platform. For the LR-perturbed multiome experiment, nuclei were processed using the 10x Genomics Chromium Next GEM Single Cell Multiome ATAC + Gene Expression workflow. Four libraries were generated: gene expression (GEX), ATAC, CRISPR, and spatial transcriptomic (ST) barcode libraries. The GEX and ST libraries were sequenced on an Illumina NovaSeq X using a 10B 100 cycle flow cell, whereas the ATAC and CRISPR libraries were sequenced on a NovaSeq X using a 1.5B 200-cycle flow cell.

### Spatial mapping and sgRNA assignment

For the first LR perturbation dataset, we used Cell Ranger v9.0.1 with default settings to quantify gene expression and assign sgRNAs from the 10x Genomics 3’ CRISPR with Feature Barcoding libraries. Gene expression reads were aligned and quantified against the 10x Genomics GRCh38-2020-A reference transcriptome, and CRISPR Guide Capture libraries were processed using the corresponding CRISPR feature reference. For the kinase perturbation dataset, which consisted of nine physical slides distributed across 13 sequencing runs, we used Cell Ranger v9.0.1 with a modified run level processing workflow. Because the kinase experiment was sequenced using the Ultima Genomics platform, Ultima unaligned CRAM files were first converted to paired end FASTQs. Each of the 13 sequencing runs was then processed independently using a sample specific Cell Ranger libraries CSV specifying one Gene Expression library and one CRISPR Guide Capture library. Gene expression reads were aligned and quantified against the 10x Genomics GRCh38-2020-A reference transcriptome using cellranger count with --libraries, --feature-ref, --localcores 16, --localmem 120, and --create-bam=false. Feature Barcode analysis was enabled by declaring the CRISPR library as CRISPR Guide Capture and supplying a CRISPR feature reference containing 1,505 guide features. The CRISPR feature reference used the downstream tethered pattern (BC)GTTTAAGAGCTAAGCTGGAA, which identifies the 20-bp protospacer immediately upstream of the conserved scaffold sequence. Cell Ranger v9.0.1 generated protospacer calls and protospacer specific UMI thresholds for all 13 runs. These thresholds were used to determine confident guide assignments while retaining cells with multiple guide assignments. For spatial processing of the kinase dataset, each of the 13 sequencing runs was processed independently through the Trekker spatial workflow on Latch Bio using the corresponding spatial transcriptomic (ST) FASTQs and matched Cell Ranger outputs. Spatial positions were assigned by extracting single-nucleus and spatial barcodes from the Trekker sequencing libraries and matching spatial barcode information to individual nuclei. The resulting Trekker output for each run was an AnnData object containing the gene expression matrix, cell-level metadata, and spatial coordinates. The Trekker-derived AnnData object was used as the base object for each run, and Cell Ranger CRISPR guide calls were incorporated from crispr_analysis/protospacer_calls_per_cell.csv and crispr_analysis/protospacer_umi_thresholds.csv. Multi-guide assignments were expanded into individual cell protospacer pairs, and guide calls were retained when the guide UMI count was greater than or equal to the Cell Ranger estimated UMI threshold for the corresponding protospacer. Cell-by-protospacer UMI and binary detection matrices were generated, and protospacer level calls were additionally collapsed to their corresponding target genes to generate a binary cell by target perturbation matrix. The 13 run level objects were subsequently combined into nine physical slide level AnnData objects. For slides represented by two sequencing runs, cell barcodes were prefixed with their Cell Ranger run identifier to prevent barcode overlap, and run and physical slide identifiers were retained as metadata. Guide matrices were aligned to the union of protospacers across runs and gene level perturbation matrices were aligned to the union of perturbed target genes, with absent features represented as zero. Run level objects belonging to the same physical slide were then concatenated along the cell axis, preserving gene expression, spatial coordinates, guide assignments, and run provenance. For the LR perturbation multiome dataset, we used an analysis workflow adapted from the CAT-ATAC framework described by Shevade et al.^38^, which extends the 10x Genomics Multiome assay to simultaneously capture gene expression, chromatin accessibility, and CRISPR perturbation identity from individual cells. The gene expression and chromatin-accessibility libraries were processed using Cell Ranger ARC v2.2.0, and CRISPR guide information and Trekker-derived spatial coordinates were integrated with the RNA and ATAC modalities to generate a multimodal MuData object. The resulting merged dataset contained 26,148 nuclei with aligned RNA, ATAC, CRISPR-guide, and spatial information.

### Quantifying Transposable Elements Activity from Single-cell Multiome Data

To assess transposable element (TE) activity, we developed an initial pipeline to quantify TE accessibility from single-cell ATAC and TE expression from single-cell RNA data. Because ATAC-seq data contains reads that span the junction between TEs and flanking sequences, we quantified the accessibility of individual TE insertions using junction ATAC-seq reads. After implementing quality control, we utilized bedtools to intersect the quality read alignments with the annotated mouse TE location information in DFam^39^. We then summed the counts of individual TE insertions to obtain total counts for each TE family. To obtain cell-level metrics, we first summed the fragment counts of the same cell barcode. We then normalized the cell-by-TE matrices by total sequence length of TE family insertions and by total fragments per cell to obtain counts per kilobase per million reads. Afterwards, we log-transformed and scaled normalized counts. In contrast to ATAC data, RNA data generally only contains the transcribed elements but not flanking sequences. Therefore, in order to quantify the TE-activity in RNA data we first created a custom transcriptome reference using TE sequences for each TE family from the same DFam annotation. Then we used kb-python^40,41^ to pseudoalign the RNA reads to the custom TE reference and quantify the expression per cell for each of the TE families. Note that we did not attempt to quantify expression of individual elements from RNA data due to the high level of sequence duplication among elements within a family and lack of junction reads to distinguish them. As with the ATAC counts, we normalized the TE family expression counts by TE family length and by cell library size, followed by log-transformation and scaling. Of note, in our current work, we only focused on long and short interspersed nuclear elements (LINE and SINE, respectively), and long terminal repeat (LTR) families. For both RNA and ATAC data, we filtered out low-count cells in all datasets.

### Preprocessing and clustering

For multiome quality control, a common cell filter was applied across RNA, ATAC, CRISPR, and spatial modalities. Cells were retained if they had a mitochondrial transcript fraction ≤20%, between 1,000 and 40,000 total RNA UMIs, a nucleosome signal ≤2, and TSS enrichment ≥1, resulting in 17,043 retained nuclei. RNA counts were normalized to 10,000 transcripts per cell and log1p-transformed. Genes detected in fewer than five cells were removed, and the top 3,000 highly variable genes were selected using the Seurat v3 variance model. S-phase and G2/M cell-cycle scores were calculated and regressed from the highly variable gene matrix before scaling. PCA was performed using 50 components, and a nearest-neighbor graph was constructed with 30 neighbors using the first 30 principal components. UMAP was calculated with min_dist=0.3 and spread=1.0, and Leiden clustering was evaluated at resolutions 0.2, 0.4, 0.6, 0.8, and 1.0, with resolution 0.6 used for the working RNA cluster assignments. For the 3’ LR and kinase datasets, we selected the top 2,000 highly variable genes, performed principal component analysis, constructed a neighbor graph, and performed community detection using the Leiden algorithm in Scanpy version 1.12.3. Leiden clustering was performed first at the default resolution and subsequently at resolutions 0.02, 0.5, and 2.0. The default resolution clustering was used for cell-type annotation. We then used marker gene expression to manually annotate each cluster according to the cell types reported in Rhodes 2022^16^. For the 3’ LR and kinase datasets, spatial artifacts were removed prior to assigning cells to individual EBs. Manually identified debris-filled regions were first excluded using hand-drawn bounding boxes in spatial x/y coordinates. A radius-neighbor connectivity graph with a radius of 150 was then constructed, and spatially isolated connected components containing three or fewer cells were removed. The remaining spatially positioned cells from each slide were clustered using DBSCAN with eps=50 and min_samples=2 to identify candidate EB clusters. Raw clusters that appeared to contain multiple EBs were re-clustered using finer DBSCAN parameters with eps=25–35 and min_samples=2. Clusters were then manually merged and assigned to biological EBs by visually comparing spatial cluster assignments with the corresponding H&E stain for each slide.

For the LR perturbation multiome dataset, EB assignment was performed separately using the two-dimensional Trekker spatial coordinates. The 17,043 QC-passing nuclei were over-segmented using K-means clustering into 200 small spatial regions using k-means++ initialization, 20 initializations, and a fixed random state of 0. The numbered spatial regions were manually inspected and spatially adjacent regions were aggregated into individual EBs, resulting in 51 assigned EBs, with regions that could not be confidently assigned retained as unassigned. RNA clusters were manually annotated using UMAP expression patterns and cluster-level expression of curated marker panels representing early ectoderm, mesoderm, neural crest, endoderm, pluripotent cells, neurons, and endothelial cells^16^. These marker-based annotations were used to assign the working RNA clusters to their corresponding cell-type identities.

### Clonal expansion analysis

To identify spatially adjacent cells that likely developed from clonal expansion during EB formation, we built a spatial graph of cells sharing identical sgRNA assignments and identified connected components. Specifically, each nucleus with an assigned spatial position is a vertex in the graph, and two vertices are connected with an edge if their spatial positions are within 100 microns of each other and identical sgRNAs were detected in the cells (including the exact combination of sgRNAs in case of multiple detections). We then identified connected components in this graph, interpreting each as a likely clonal group of cells that developed from local proliferation of a transduced cell that passed its integrated sgRNA combination to its daughter cells.

### Identifying distance-dependent non-autonomous responses to ligand knockdown

CytoSignal ligand–receptor (LR) scores were matched to nuclei in the spatial multiome dataset. Direct KD–LR tests were defined as interactions in which the perturbed gene matched either the ligand or receptor component of the LR pair. Compensatory interactions were specified separately and matched to the LR table by interaction ID or LR name. For each perturbation, cells carrying a guide targeting the gene of interest were used as spatial reference cells, and cells not carrying that perturbation were treated as receiver cells. Analyses were performed within individual EBs; an EB was included when it contained at least two KD cells and at least ten non-KD cells. For each receiver cell, the distance to the nearest KD cell within the same EB was calculated, and receivers within 300 µm were pooled across eligible EBs for the dataset-level test. For each KD–LR pair, distance dependence was quantified using the Spearman correlation between receiver-cell distance and CytoSignal LR score. Statistical significance was determined by 100,000 spatial permutations. For each permutation, whole-cell perturbation labels and corresponding LR-score profiles were shuffled together across spatial positions, after which KD reference cells, receiver cells and their distances were recalculated within each EB. The positive- and negative-direction empirical *P* values were calculated as

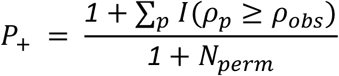

and

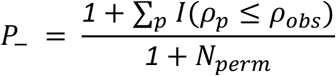

where *ρ*_*obs*_ is the observed Spearman correlation and *ρ*_*p*_ is the correlation obtained after permutation. Positive and negative one-sided *P* values were corrected separately using the Benjamini–Hochberg procedure across LR interactions tested for each KD, with FDR < 0.05 considered significant.

For visualization, receiver cells were grouped into six 50-µm intervals spanning 0–300 µm. The mean CytoSignal LR score and cell-level s.e.m. were calculated for each interval. These bins were used only to display the spatial LR profiles; the statistical test used the individual receiver-cell distances and LR scores.

#### Identifying distance-dependent epigenomic responses to ligand knockdown

Distance-dependent epigenomic responses were tested for 15 ligand perturbations across four ATAC-derived feature spaces: called ATAC peaks, fixed 50-kb SnapATAC2 bins, chromVAR motif-deviation scores and Signac ATAC gene-activity scores. For each ligand, nuclei assigned to that ligand perturbation were used as reference cells, and non-target nuclei were used as query cells. Distances to the nearest ligand-KD nucleus were calculated separately within each EB and restricted to 300 µm. Called ATAC peaks were grouped into six 50-µm intervals, whereas 50-kb bins, chromVAR scores and gene-activity features were grouped into twelve 25-µm intervals. For count-based modalities, features detected in more than 5% of eligible cells were retained. Counts were summed across cells within each distance interval and normalized according to

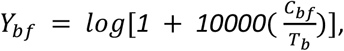

where *C*_*bf*_ is the summed count for feature *f* in distance bin *b*, and *T*_*b*_ is the summed library size of the cells in that bin. For called peaks, library size was based on total ATAC counts; 50-kb bins and gene activity used the corresponding full count matrix before feature filtering. For these count-based profiles, uncertainty was calculated using a ratio-estimator and delta-method approach that retained the covariance between feature counts and library size.

Defining

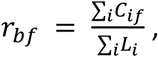

the variance of the ratio was estimated as

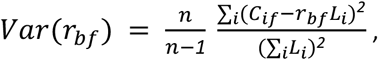

and the s.e.m. after log transformation was

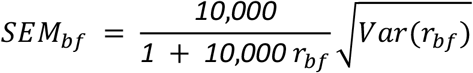

For chromVAR, the mean motif-deviation score and standard cell-level s.e.m. were calculated directly within each distance interval. Distance dependence was tested independently for each feature by comparing two zero-mean Gaussian-process models. The null model used a constant covariance kernel,

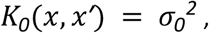

whereas the alternative model used a radial-basis-function kernel,

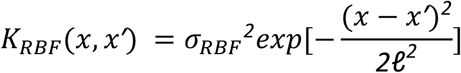

Observation-noise variance was fixed for each feature to the mean squared s.e.m. across distance bins,

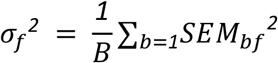

The RBF model was optimized from starting length scales of 10, 50 and 200 µm using up to 1,000 L-BFGS-B iterations, and the fit with the highest likelihood was retained. The primary analysis did not impose a minimum biological length scale. The likelihood-ratio statistic was calculated as

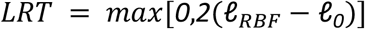

Because the constant model is approached as the RBF length scale tends to infinity, significance was evaluated using a *0*.*5χ*_*0*_^*2*^ + *0*.*5χ*_*1*_^*2*^ boundary distribution. For *LRT* > *0*,

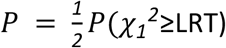

Benjamini–Hochberg correction was performed separately within each perturbation and modality, with FDR ≤ 0.05 considered significant. Significant features were classified as increasing or decreasing with distance according to the sign of the Spearman correlation between their distance-bin means and distance.

### Ligand-driven enhancer-gene regulatory network analysis

We used CisTarget and GRNBoost2^20^ to identify chromatin accessibility peaks and genes that are likely regulated by each transcription factor in a database of binding motifs for 1,553 human transcription factors. Briefly, these approaches use a combination of motif occurrence and co-expression (or expression-accessibility correlation) to link TFs to candidate downstream peak and gene targets. We then used the results of the distance-dependent ligand knockdown analysis described in the previous section to identify transcription factors whose expression increases or decreases with distance from cells with a particular ligand knocked down. These ligand-driven transcription factors represent candidates for regulatory effectors of signal transduction pathways mediated by ligand signaling. We then built a ligand-driven enhancer-gene regulatory network by connecting a ligand to a TF with a positive edge if the TF showed an expression trend increasing with distance from knockdown cells, and with a negative edge if the TF showed an expression trend decreasing with distance from knockdown cells.

### Detecting context-dependent responses to receptor and kinase knockdown

We used CytoSignal to estimate the local ligand–receptor signaling environment surrounding each cell. We calculated LR scores independently at diffusion radii of 100, 200, and 400 µm. Ligand expression was spatially diffused over the corresponding radius, whereas receptor expression was retained at its observed value. LR activity scores were then calculated using the CytoSignal diffusion-dependent interaction database and smoothed over the spatial graph. We first performed an elastic-net screen to prioritize a smaller set of LR scores for each perturbation target and spatial radius. We represented the transcriptome using 30 RNA principal components derived from up to 8,000 highly variable genes. Models were fit separately for each perturbation target and spatial radius using cells assigned to the corresponding knockdown and cells assigned to non-targeting controls. For receptor knockdowns, the elastic-net model included the knockdown indicator, LR-score main effects, and knockdown-by-LR interaction terms. LR pairs were ranked primarily according to the magnitude of their knockdown-by-LR interaction coefficients across the RNA principal components. For kinase knockdowns, due to the substantially larger number of kinase perturbations, we therefore used a lighter first-stage screen in which the LR scores were used to predict the RNA principal components within cells carrying the kinase knockdown and non-targeting controls. LR pairs were ranked according to the aggregate magnitude of their coefficients across the RNA principal components.

We then carried only the top-ranked LR scores forward to gene-level regression to identify downstream genes whose knockdown responses varied with the local LR signaling environment. For each selected knockdown target, LR pair, spatial radius, and expressed gene, we fit the following ordinary least-squares model:

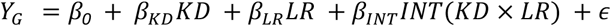

Here, *Y*_*G*_ is the expression of gene (g), *KD* indicates whether the cell carried the target-specific guide or a non-targeting control guide, and *LR* is the standardized LR score. Genes detected in fewer than 5% of the cells included in a model were excluded. The interaction coefficient, *β*_*INT*_*INT*, was used to identify context-dependent knockdown responses.

We used two-sided (t)-tests to evaluate the knockdown, LR-score, and knockdown-by-LR interaction coefficients. For each coefficient family, raw (p)-values were pooled across all perturbation targets, selected LR pairs, genes, and spatial radii, and the Benjamini–Hochberg procedure was applied to control the project-wide false discovery rate. Context-dependent responses were identified based on the knockdown-by-LR interaction term, using a project-wide FDR threshold of 0.05. Significant responses were subsequently ranked according to the number of genes with significant interaction effects, while the magnitude and direction of the interaction coefficients were used to interpret individual target genes.

To visualize representative examples of the context-dependent responses, we generated spatial maps for selected perturbation–LR–gene combinations with a project-wide interaction FDR below 0.05. For each combination, EBs were evaluated separately. We prioritized examples in which gene expression showed a clear association with the LR score within knockdown cells and in which this association differed from that observed in control cells. The direction of the EB-level slope difference was also compared with the sign of the interaction coefficient from the project-wide regression. The highest-ranked EBs were used as representative spatial examples in the map panels.

### Identifying trends in spatial locations of kinase perturbations

We computed two orthogonal spatial axes for each embryoid body (EB) by applying principal component analysis (PCA) to the two-dimensional spatial coordinates of all cells within the EB. We retained EBs with at least 50 cells and normalized each axis projection to the range [−1, 1] by dividing by the maximum absolute value. The first principal component defined the major axis and the second defined the minor axis.

The sign and orientation of the spatial PCs are arbitrary across EBs. To assign a consistent biological orientation to each axis, we correlated gene expression with axis position using Spearman’s rank correlation, requiring the marker gene to be expressed in at least 10% of cells within the EB. We found GATA4 and GRID2 correlated with the major and minor axes, respectively. If the Spearman correlation was negative, we flipped the axis sign so that the positive direction corresponded to higher marker expression (e.g., GATA4+ for the major axis, GRID2+ for the minor axis). We determined axis orientations once and held them fixed across all subsequent analyses.

To test whether kinase knockdown cells occupied non-random positions along each oriented axis, we used a within-EB permutation test. For each kinase with at least 20 cells across all oriented EBs, we computed a weighted mean signed axis position, weighting per-EB means by the number of kinase-targeted cells in each EB. Cells carrying guides targeting multiple kinases contributed to each kinase independently. We generated a null distribution by shuffling complete guide-label strings among all cells within each EB (preserving the within-EB guide composition) 1,000 times. We computed two-sided empirical p-values as the fraction of permutations for which |stat_perm − mean_null| ≥ |stat_obs − mean_null|, and derived z-scores as (stat_obs − mean_null) / std_null. We corrected for multiple testing using the Benjamini–Hochberg procedure, applied separately for each axis, excluding non-targeting control (NTC) cells from the correction.

As an alternative to single-gene axis orientation, we developed a multi-gene consensus method. For each EB, we computed the Spearman correlation of each of the top 5,000 highly variable genes (selected by coefficient of variation, requiring expression in at least 1% of cells) with the major and minor axis positions. We Fisher-transformed the correlations (z = arctanh(ρ)) and combined them across EBs using Stouffer’s weighted meta-analysis, with weights proportional to √(n − 3) where n is the number of cells per EB. We initialized axis orientations using the single-gene markers described above, then iteratively refined them: at each iteration, we identified the top 50 genes by absolute meta-z, computed a weighted vote for each EB based on the concordance of its per-gene correlations with the consensus signs, and flipped EBs with a negative vote. We repeated this procedure until no further flips occurred (convergence in 7 iterations). The multi-gene orientations agreed with the single-marker orientations for 68% of EBs on the major axis and 73% on the minor axis.

### Modeling spatial perturbation response with graph neural networks

#### Spatial graph and model inputs

We developed a modality-specific spatial-residual graph attention network (GAT) to model cell-non-autonomous perturbation effects. Each nucleus was represented as a node, and spatially neighboring nuclei within the same embryoid body (EB) were connected using a *k*-nearest-neighbor graph. Self-edges and edges between different EBs were excluded. The input for each nucleus consisted of its binary CRISPR guide assignments and one-hot encoded cell type. RNA expression and ATAC accessibility were prediction targets and were not used as input features. RNA and ATAC were modeled with separate branches, allowing their spatial graph parameters to be tuned independently. For each training run, one EB was held out for testing and a separate EB was used for validation, for a total of ten held-out test EBs. RNA and ATAC target features were selected using training cells only. For RNA, counts were normalized to 10,000 counts per cell, log transformed and ranked by variance, and the top 1,000 genes were retained. For ATAC, the top 1,000 features were selected using Poisson deviance calculated from raw counts.

#### GAT architecture

For each modality, a non-spatial multilayer perceptron (MLP) was first trained using only the guide and cell-type features of the nucleus itself. The MLP included direct guide and cell-type effects together with a nonlinear encoder comprising sequential 128- and 64-dimensional hidden layers. The spatial model used the trained MLP as its non-spatial baseline and learned an additional spatial correction from neighboring cells. The input for each cell included its own guide and cell-type identities, the guide and cell-type identities of its spatial neighbors, and the distances between the cell and its neighbors. Spatial distance between cells was also included as an edge feature. A single GATv2 layer with four attention heads of 32 dimensions each generated a 128-dimensional spatial representation. The model also included a distance-weighted summary of neighboring guide identities, neighboring cell types, number of neighbors and neighbor distances.

The spatial representation was passed through a gating layer and a correction decoder. The final spatial correction layer was initialized to zero, so the GAT initially produced the same prediction as the trained MLP. During GAT training, the MLP parameters were frozen and only the spatial components were trained. Thus, the final prediction was

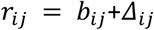

where *b*_*ij*_ is the non-spatial MLP prediction and Δ_*ij*_ is the spatial correction learned from neighboring cells. RNA and ATAC counts were modeled with a negative-binomial distribution. The predicted mean count for cell *i* and feature *j* was

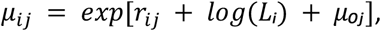

where *r*_*ij*_ is the model output, *L*_*i*_ is the cell-specific non-target library-size offset and *μ*_*oj*_ is the baseline feature rate estimated from the training cells. A feature-specific negative-binomial dispersion parameter was also learned. The loss for each modality combined negative-binomial negative log-likelihood with log-transformed mean-squared error, Pearson correlation and normalized mean-squared error:

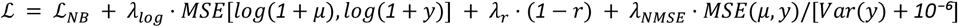

Loss weights and spatial graph parameters were selected using the validation EB separately for RNA and ATAC. Models were trained with AdamW, a learning rate of 0.003, weight decay of *10*^*−5*^, gradient clipping at 1.0 and early stopping with a patience of 25 epochs. Final models were trained with three random seeds (17, 42 and 89), and their predictions were averaged to generate the final three-seed ensemble prediction.

#### Counterfactual inference of non-autonomous perturbation effects

The trained GATs were then used for counterfactual inference without retraining the model. For each receiver cell, the original neighborhood was first used to generate a baseline RNA and ATAC prediction. Guide identities were then changed only in neighboring sender cells *in silico*, and the trained GAT was rerun. The receiver cell itself was never edited. For guide *A* → *B* comparisons, receiver cells carrying either guide *A* or guide *B* were excluded. The counterfactual effect for each predicted feature was calculated as

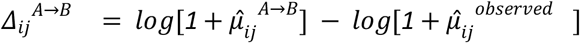

Where 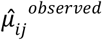 is the prediction using the original neighborhood and 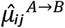 is the prediction after replacing guide *A* with guide *B* in neighboring sender cells. A→B and B→A were evaluated separately. Predictions were averaged across the three trained GAT seeds. For the guide-pair analysis, all ordered pairs among the ten selected non-control guides were tested.

For RNA visualization, genes were selected by expression variability rather than by the size of their GAT effect. Mitochondrial, ribosomal and common housekeeping genes were removed, and the 100 highest-variance informative genes were retained. Guide-pair RNA and ATAC effects were summarized using the mean absolute log1p counterfactual change across receiver cells.

#### Distance-resolved counterfactual analysis

To measure how non-autonomous perturbation effects changed with distance, neighboring perturbation-positive cells were divided into distinct spatial distance bands. Within each band, the perturbation guide in neighboring sender cells was replaced *in silico* with the non-targeting control guide, while the receiver cell remained unchanged. RNA and ATAC predictions before and after the edit were compared separately for each distance band. Distance bands were generated separately for each run from the range of neighbor distances in the fitted graphs, using up to four equal-width bands with a minimum width of 10 µm. Because the exact physical boundaries differed slightly between runs, results were combined across runs using the matching within-run band number rather than treating the different physical boundaries as separate distances. The displayed distance for each common band was the mean midpoint across contributing runs. For each perturbation and distance band, effects were summarized using 1,000 EB-level bootstrap resamples. The bootstrap distribution was used to calculate 95% confidence intervals and standard errors. Two-sided 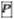 values were calculated from the bootstrap standard error and corrected using the Benjamini–Hochberg method, with FDR < 0.05 considered significant.

#### Spatial effective radius

For each receiver cell, the distance-resolved effects were also summarized as an effect-weighted spatial radius. The effective radius was calculated as

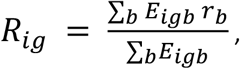

where *E*_*igb*_ is the mean absolute GAT counterfactual effect for receiver cell *i*, perturbation *g* and distance band *b*, and *r*_*b*_ is the midpoint of that distance band. Distance bands with stronger predicted effects therefore contributed more strongly to the effective-radius estimate. For RNA, the radius was calculated using the same variance-selected informative genes used in the guide-pair analysis; ATAC used the modeled ATAC features. In the spatial maps, receiver-cell color represents the weighted effective radius in micrometers and point size represents the strength of the predicted effect. These values describe the spatial range of the GAT-predicted non-autonomous effect, rather than a physical boundary of signaling.

## Data and Code Availability

Raw and processed sequencing data are available in the Gene Expression Omnibus (GSE342761) Code for reproducing the analyses in the paper is available at https://github.com/welch-lab/perturb-tags/

## Author Contributions

J.D.W. conceived the study. J.G. designed and performed the experiments. J.G., with help from K.L., generated the data. J.G., Y.W., D.C., and J.D.W. designed the graph neural network approach. J.G. implemented the graph neural network. J.G., J.D.W., and L.L. performed data analyses and wrote the manuscript. L.L., L.L., G.G., M.W., Y.Z., S.J., and K.L. contributed to data analysis.

## Acknowledgments

We thank the University of Michigan Advanced Genomics Core for their invaluable assistance in generating the Trekker data, including tissue sectioning, 10X Genomics library preparation, and Ultima sequencing. We also gratefully acknowledge support from the Count on Us program sponsored by Ultima. This study was supported by the University of Michigan Bold Challenges program, National Institutes of Health grant R01HG010883 to J.D.W., and J.G. was supported by NIH training grant T32HG000040.

